# Expression of Most Retrotransposons in Human Blood Correlates with Biological Aging

**DOI:** 10.1101/2024.02.09.579582

**Authors:** Yi-Ting Tsai, Nogayhan Seymen, I. Richard Thompson, Xinchen Zou, Warisha Mumtaz, Sila Gerlevik, Ghulam J. Mufti, Mohammad M. Karimi

## Abstract

Retrotransposons (RTEs) have been postulated to reactivate with age and contribute to aging through activated innate immune response and inflammation. Here, we analyzed the relationship between RTE expression and aging using published transcriptomic and methylomic datasets of human blood. Despite no observed correlation between RTE activity and chronological age, the expression of most RTE classes and families except short interspersed nuclear elements (SINEs) correlated with biological age-associated gene signature scores. Strikingly, we found that the expression of SINEs was linked to upregulated DNA repair pathways in multiple cohorts. We also observed DNA hypomethylation with aging and significant increase in RTE expression level in hypomethylated RTEs except for SINEs. Additionally, our single-cell transcriptomic analysis suggested a role for plasma cells in aging mediated by RTEs. Altogether, our multi-omics analysis of large human cohorts highlights the role of RTEs in biological aging and suggests possible mechanisms and cell populations for future investigations.

## Introduction

Transposable elements (TEs) are genetic elements that can move within the genome and are categorized into DNA transposons and retrotransposons (RTEs), which depend on cDNA intermediates to function. RTEs include three classes: endogenous retrovirus long terminal repeats (LTRs), and long and short interspersed nuclear elements, known as LINEs and SINEs, respectively. LINEs and SINEs employ target-primed reverse transcription for genome integration, with SINEs depending on the proteins encoded by LINEs^1^. Although most of the RTE sequences are dormant in the host genome, they continue to play vital roles in human evolution and physiology^2,3^.

RTEs are silenced through heterochromatinization and DNA methylation in the early developmental stages as part of the host defense mechanism^3,4^. However, increased RTE activity that occurs with aging can lead to genome instability and activation of DNA damage pathways^4–7^. Additionally, accumulation of cytoplasmic RTE cDNAs detected in aging organisms and senescent cells can activate type I interferon (IFN-I) response and inflammation^3,4,8–10^, that contributes to inflammaging^11^. Furthermore, chromatin remodeling and cellular senescence also contribute partially to the RTE reactivation^10^. Cellular senescence is a non-proliferative state elicited by stress factors such as DNA damage and is closely intertwined with inflammation and aging^12^. Through the expression of senescence associated secretory phenotype (SASP), senescent cells promote inflammation and senescence in other cells, including immune cells, leading to a compromised immune system and a vicious cycle of inflammation^13^. Understanding the relationship between RTE expression and SASP, cellular senescence, and inflammaging could offer important insights into strategies against aging and age-related diseases (ARDs).

Recent studies have documented the relationship between RTE activation and biological age-related (BAR) events, but comprehensive and large-scale studies are lacking, mainly due to the paucity of RNA-seq data for large non-cancerous human cohorts. Most of the existing transcriptomics datasets are microarray-based, where the computational methods to analyze repetitive elements have not been sufficiently developed. By overlapping Illumina microarray expression and methylation probe locations to RTE locations in RepeatMasker^14^, we were able to identify sufficient number of probes to calculate the expression and methylation levels of RTE classes and families. Building on this methodology, we explored how RTE expression contributes to biological aging using publicly available transcriptomics microarray data derived from human blood samples. More specifically, we first investigated if RTE expression was correlated with chronological age, and then we analyzed the relationship between RTE and BAR events including cellular senescence, inflammation, and IFN-I response with published gene signatures. Using microarray methylomic data, we also investigated the DNA methylation level of RTEs in blood samples of multiple non-cancerous human cohorts and examined the relation between DNA methylation and RTE expression and aging. Furthermore, with annotated single-cell transcriptomic data of the peripheral blood mononuclear cells (PBMCs) of 21 healthy human samples from a recent study of aging immunity^15^, we identified cells that could be implicated in the process of RTE reactivation and aging. We also validated our microarray result with a publicly available bulk RNA-seq PBMC data from healthy individuals that was recently published in a new study of aging^16^. Lastly, the single-cell transcriptomic data of supercentenarians^17^ was analyzed to examine the interplay between RTE expression and aging as a factor of aging and longevity (**Fig. 1**).

**Figure 1.**
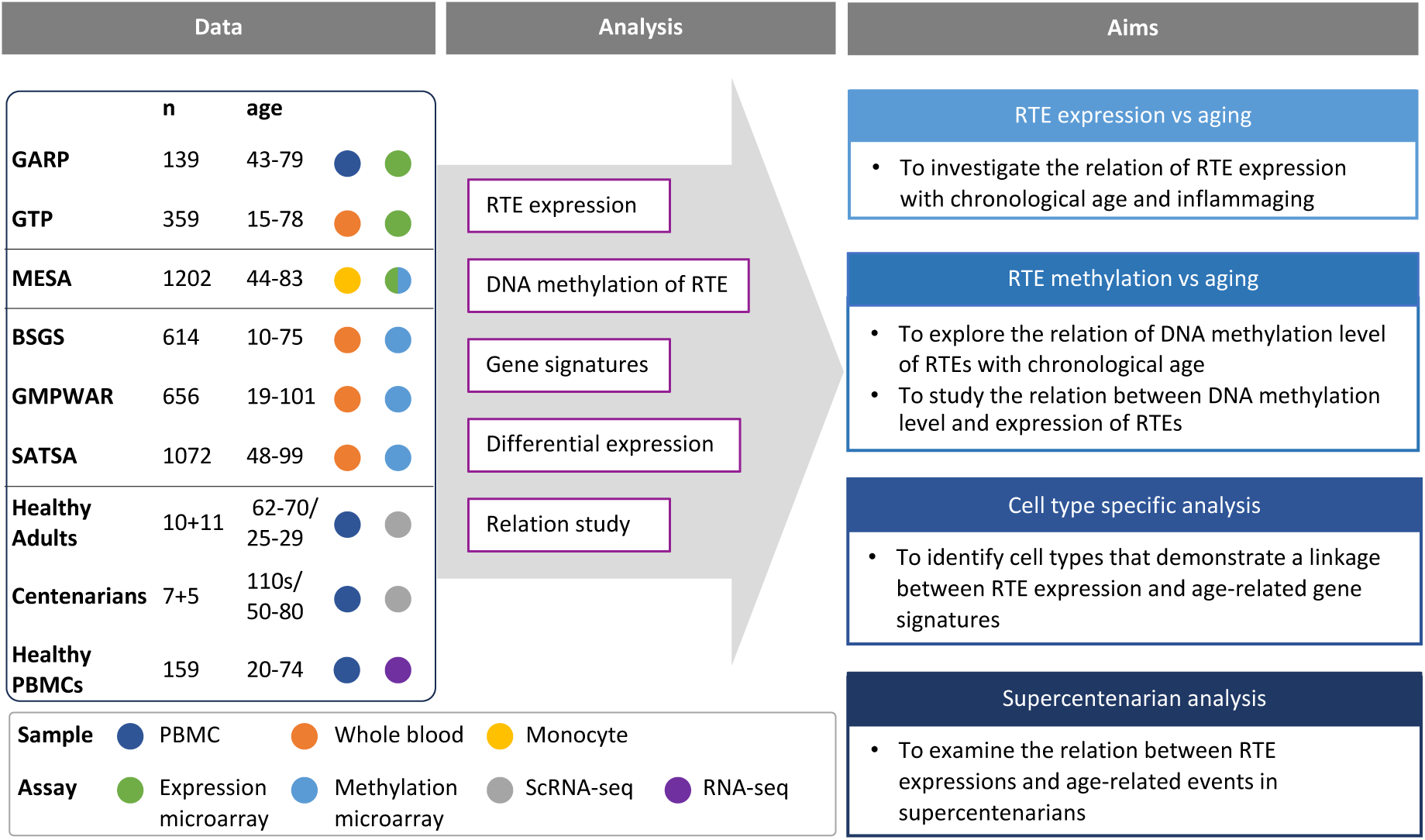
Conceptual framework and the study design. We collected published datasets of human blood samples for gene expression, DNA methylation, and single-cell transcriptomic data. The analysis aimed to study the relation between the expression and DNA methylation of RTEs versus chronological and biological aging in large human cohorts. The single-cell transcriptomic datasets were employed for cell type-specific analysis of RTEs in PBMC to identify the relation between RTE expression and aging events for annotated cell types within old versus young PBMC samples.

## Results

### Analyzing DNA methylation and expression levels of RTEs using microarray data

We collected three published microarray datasets from large-scale human studies, including the peripheral monocyte samples from Multi-Ethnic Study of Atherosclerosis^18^ (MESA, aged 44 to 83, n=1202), the whole blood (WB) samples from Grady Trauma Project^19,20^ (GTP, aged 15 to 77, n=359), and PBMC samples from Genetics, Osteoarthritis and Progression^21^ (GARP, aged 43 to 79, n=139), which are all non-cancerous human samples (**Supplementary Table 1** and **Supplementary Information**). The studies were conducted using either Illumina HumanHT-12 V3 or Illumina HumanHT-12 V4 expression microarray kits. After comparing the probes of the two microarray versions, we adopted the more comprehensive probe list of V4, which contains the full intersection of the two lists. To quantify RTE expression, we mapped the microarray probe locations to RTE locations in RepeatMasker to extract the list of noncoding (intergenic or intronic) probes that cover the RTE regions. We included three main RTE classes: (1) the LINE class, which encompasses the L1 and L2 families; (2) the SINE class, with Alu and MIR as the two main families; and (3) the LTR class which comprises the ERV1, ERVL, ERVL-MaLR, and ERVK families. Most of the RTE-covering probes available on Illumina HumanHT-12 V4 are present in MESA and GARP, while fewer are available in GTP (**Supplementary Fig. 1a**, **Supplementary Tables 2-4**).

Four methylation datasets were analysed, including MESA, Swedish Adoption/Twin Study of Aging^22^ (SATSA, aged 48 to 98, n=1072), Brisbane Systems Genetics Study^23^ (BSGS, aged 10 to 75, n=862), and Genome-wide Methylation Profiles Reveal Quantitative Views of Human Aging Rates^24^ (GMPWAR, aged 19 to 101, n=656). SATSA, BSGS, and GMPWAR include genome-wide DNA methylation of WB samples produced by Illumina Infinium 450k array (**Supplementary Table 1** and **Supplementary Information**). The DNA methylation probes from the Illumina Infinium 450k array kit were aligned to the locations in RepeatMasker to identify the probes overlapping the RTE regions (**Supplementary Table 5**). More than 90% of the RTE-covering probes are present in MESA and GMPWAR, while fewer, but more than half, are available in SATA and BSGS datasets (**Supplementary Fig. 1b**).

### The chronological age is not linked with RTE expression

Firstly, we examined the relationship between RTE expression and chronological age in the MESA, GARP, and GTP cohorts. The ages of the individuals enrolled in these studies range from 15 to 83 years, with the mean being 70.2 (MESA), 60.1 (GAPR), and 42.5 (GTP) (**Supplementary Table 1**). Strikingly, no significant relationship (*P* value < 0.05) is found between chronological age and expressions of the three RTE classes across the datasets (**Fig. 2a**). Similarly, apart from a weak correlation observed between a few RTE families and chronological age, we could not identify any strong relation between chronological age and expression of RTE families across the three cohorts (**Supplementary Fig. 2a**).

**Figure 2.**
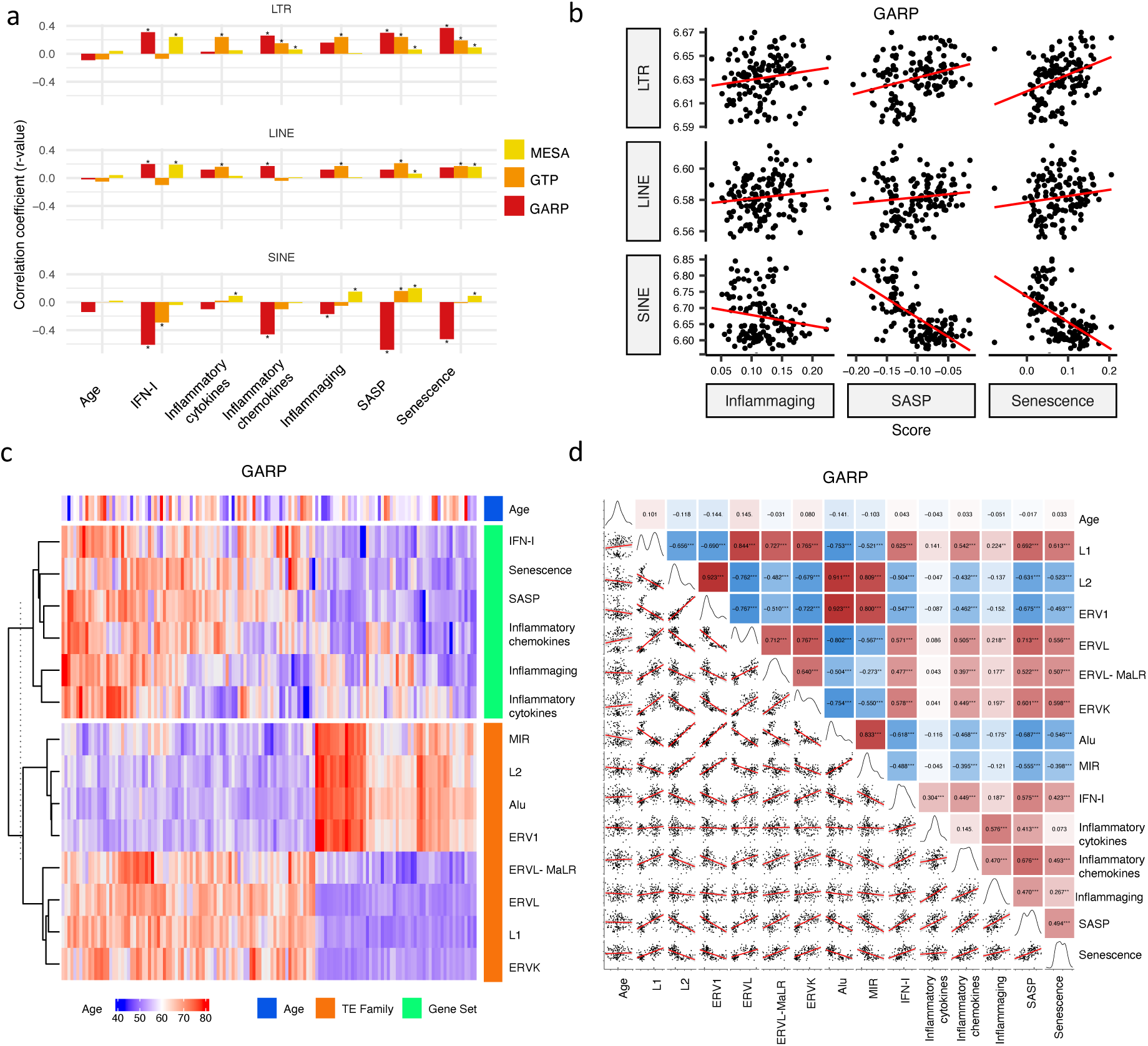
Correlation analysis between RTE expression, chronological age, and age-associated gene signature scores. **a**, No correlation between RTE expression and chronological age versus positive correlations between BAR gene signature scores and LINE and LTR expressions. Pair-wise correlation coefficients were calculated between the expression of different RTE classes (LTR, LINE & SINE) and chronological age and six BAR gene signature scores in monocytes (MESA), PBMCs (GARP), and the WB (GTP). **b**, Scatter plots displaying a positive correlation between LINE and LTR expressions and inflammaging, SASP, and senescence gene signature scores in PBMCs. **c**, Different families of RTEs were divided into two major groups based on their correlation and inverse correlation with BAR gene signature scores in PBMC samples. **d**, Correlation matrix depicting all pair-wise combinations to identify the correlation between chronological age, RTE family expressions, and six age-associated signature scores in PBMCs. ** *P* ≤ 0.01, *** *P* ≤ 0.001, Pearson’s correlation. MESA, n=1202; GARP, n=139; GTP, n=359.

A similar correlation analysis was carried out in a scRNA-seq data of the PBMCs of 21 non-obese healthy men annotated into 25 cell types by Mogilenko et al.^15^ (**Supplementary Table 6** and **online methods**). When comparing the RTE expression of the young (aged 25 to 29, n=11) group with the old (aged 62 to 70, n=10) group, we could not find any significant difference or trend in any cell types between the two groups (**Supplementary Figs. 3-5**).

To investigate whether our observation is consistent in whole transcriptomic data, we conducted correlation analysis on a set of RNA-seq data on healthy human PBMC^16^ (aged 20 to 74, n=159). We did not observe any statistically significant correlation between chronological age and expression of RTE classes or families (**Supplementary Fig. 6**). Overall, RTE expression did not correlate with chronological aging across the microarray, scRNA-seq, and RNA-seq datasets.

### RTE expression positively correlates with BAR gene signature scores except for SINEs

In mammals, aging is characterized by an intricate network of inflammation, IFN-I signaling, and cellular senescence^3^. To measure the level of biological aging from the transcriptomic data, we focused on six gene sets retrieved from widely cited studies: IFN-I^25^, inflammatory cytokines^26^, inflammatory chemokines^27^, inflammaging^28^, SASP^29^, and cellular senescence^30^ (**Supplementary Table 7**). The scores of these six age-associated gene sets were calculated for each individual in the MESA, GTP, and GARP cohorts by applying the *singscore* package in Bioconductor^31^ (**online methods**). Except for GTP, in which 12 genes were missing from the gene sets in total (**Supplementary Table 8**), MESA and GARP microarray datasets had the expression of all the genes in the gene sets.

We analyzed the relation between RTE expression and the six BAR gene signature scores. Positive correlations were observed between the expressions of LINEs and LTRs and BAR gene signature scores across three datasets, while inverse correlation was present between the expression of SINEs and these scores in the PBMC samples from the GARP cohort (**Fig. 2a** and **b**). The discrepant pattern of LTRs and LINEs versus SINEs in PBMC is more pronounced in the correlation analysis between the expression of RTE families and the BAR gene sets (**Fig. 2c** and **d**). Interestingly, the expressive patterns of RTE families are divided into two groups that also reflect their positive or inverse correlations with BAR signature scores in the GARP cohort. In the GARP cohort, the strongest positive correlation with BAR scores was observed in L1, ERVK, and ERVL. By contrast, the expressions of Alu and MIR of SINE, L2, and ERV1 were inversely correlated with BAR signature scores in the GARP cohort (**Fig. 2c** and **d**). This pattern was also observed in the MESA peripheral monocyte samples for L1, ERVL, and ERVK, but not other RTE families (**Supplementary Fig.2a and c**). The WB GTP cohort showed divergent result to the other two cohorts perhaps because of the absence of certain genes in the six BAR gene sets or lower number of TE probes in GTP (**Supplementary Fig. 1b, Fig.2a-b** and **Supplementary Table 8**).

To study the association between RTE expression and BAR gene signatures, we also split the samples in each cohort based on quartiles of RTE expression into high, low, and medium (top 25%, bottom 25%, and middle 50%) groups and compared the BAR gene signature scores across these three groups. Across the datasets, there is an overall trend of increased scores of BAR gene signatures in high versus low LINE and LTR expression groups (**Supplementary Fig. 8**). In fact, amongst all gene sets, we found the most significant increase for SASP and senescence and to a lesser extent for inflammaging signatures. We observed higher SASP, inflammatory cytokine, senescence, and inflammaging in the peripheral monocytes (MESA) in high SINE- and Alu-expressing groups (**Supplementary Fig. 9**). However, in the PBMC samples of the GARP cohorts, groups of high SINE and Alu expressions demonstrate lower expressions of SASP, inflammatory chemokine, senescence, and inflammaging, that reflects the inverse correlation we observed before (**Fig. 2c**). Interestingly, when we examined RNA-seq PBMC data, we found that there is a significant correlation between IFN-I signature score and expression of LINE and LTR classes and most of their families (P-val < 0.01), but such a significant correlation did not exist between IFN-I score and SINE class and Alu family (**Supplementary Fig. 7**). There are also no specific trends for SINE expression in the WB samples from the GTP cohort (**Supplementary Fig. 9**). Taken together, SINEs display different pattern than LINEs and LTRs in terms of the association between their expressions and BAR gene signatures. This prompted us to conduct a gene set variation analysis (GSVA)^32^ to identify the pathways that are regulated in the high versus low RTE expression groups in the three cohorts (**online methods**).

### Elevated SINE expression is linked with the upregulation of the DNA repair pathways, while elevated LINE and LTR expressions are associated with the inflammatory response

We conducted GSVA for high versus low expression groups of RTE classes and families to identify the pathways that might be affected by RTE expression. We specifically focused on the pathways (gene sets) related to the inflammatory and DNA repair responses retrieved from the Molecular Signatures Database (MSigDB)^33^ with “inflammatory” and “dna_repair” as search keywords.

We found that a significant fraction of gene sets related to DNA repair were downregulated in the group of monocyte samples (MESA cohort) highly expressing LINE and LTR classes but upregulated in the monocyte and PBMC samples (MESA and GARP cohorts, respectively) highly expressing SINEs (**Fig. 3a**). At the family level, there is a unique increase in DNA repair activity in the monocytes of the cohort with high Alu expression. In the PBMC samples of the GARP dataset, DNA repair pathways are upregulated in groups highly expressing L2, ERV1, Alu, and MIR, while most of the other RTE classes demonstrate decreased DNA repair activity. In contrast, the inflammatory response is upregulated in the monocyte and PBMC samples of groups expressing LINE and LTR classes and families at high levels, and such upregulation is less common in SINE (**Fig. 3a**). These observations provide insight into the perpetuating contrast between SINE versus LINE and LTR observed in our analysis, as SINE might contribute to aging via genome instability instead of inflammation that is associated with LINE and LTR activities. Overall, our observations are consistent across MESA and GARP cohorts but less so in the GTP cohort (**Fig. 3b**), which might be attributed to less RTE-covering probes and the missing genes from the BAR gene signatures. (**Supplementary Fig. 1a, Supplementary Table 8**).

**Figure 3.**
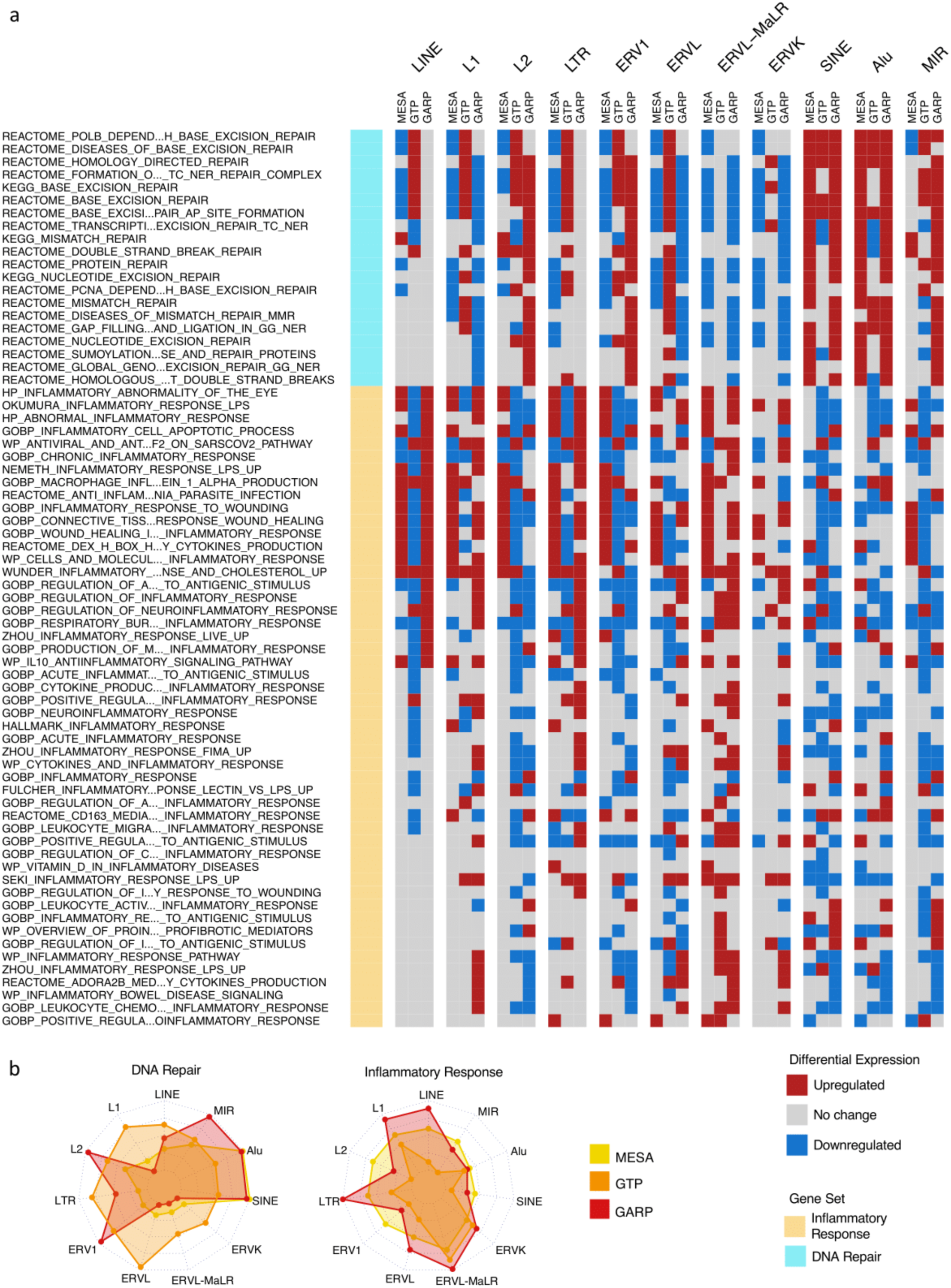
Upregulation of DNA repair vs inflammatory responses for samples with high expression of SINE vs LTR and LINE in the MESA and GARP cohorts. **a**, GSVA demonstrates increased activity of DNA repair pathways in the group of samples with high vs low SINE expression in the MESA and GARP cohorts. In contrast, the inflammatory response is upregulated in the sample groups highly expressing LINE and LTR classes and families in the MESA and GARP cohorts. The samples in each cohort were divided into low (1^st^ quartile), medium (2^nd^ and 3^rd^ quartile), and high (4^th^ quartile) expression groups based on the expression of RTE classes or families. GSVA was applied on high vs low groups for each class and family of RTEs. The threshold for differential expression is set at |logFC| > 0.1 and *P* < 0.05 (**online methods**). **b**, The Radar plot shows the difference between the number of upregulated versus downregulated gene sets related to DNA repair and inflammatory response in each cohort. While high expression of SINE and Alu is significantly associated with high number of up-regulated DNA-repair gene sets, LINE and L1 expressions are highly related to high number of activated gene sets related to inflammatory response in the MESA and GARP cohorts. This result is not highly supported by the GTP cohort, more likely due to the low number of probes in this cohort.

### DNA methylation levels of RTEs inversely correlated with the chronological age and the RTE expression except for SINE expression

The demethylation of RTEs is suggested to contribute to aging^3,5,7^. To examine the effect of RTE demethylation on chronological aging, we analyzed DNA methylation data from multiple cohorts including monocytes from the MESA cohort and the WB data from GMPWAR, BSGS, SATSA. Our analysis showed an inverse correlation between the methylation level of RTEs and chronological age across all RTE classes and families in the WB and monocytes (**Fig. 4a-b** and **Supplementary Fig. 10**). This observed trend is in line with previous reports of demethylation of RTE with age in gene-poor regions^5,22,24^.

**Figure 4.**
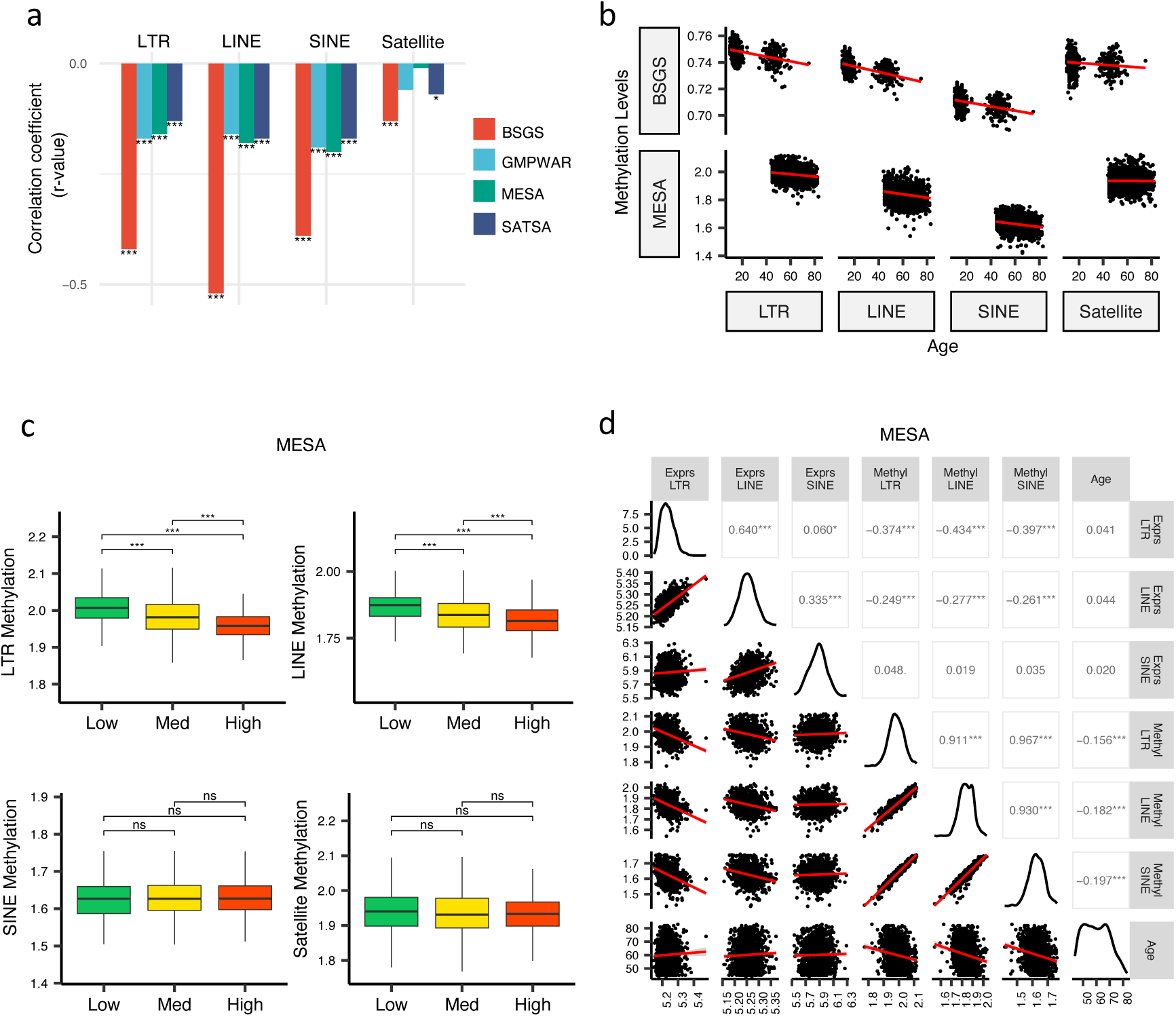
Inverse correlation of DNA methylation levels of RTEs with the chronological age and the RTE expression except for SINE expression. **a, b**, Methylation levels of RTE classes inversely correlated with chronological age in monocyte (MESA) and WB (BSGS, SATSA, and GMPWAR) samples. Satellite DNA was included as a control group. **c**, Methylation levels versus low (1st quartile), medium (2nd and 3rd quartile), and high (4th quartile) expressions of RTE classes in monocytes (MESA). Wilcoxon test; ns: not significant. **d**, Correlation matrix for RTE expressions and methylation levels, and chronological age. * *P* ≤ 0.05, **** *P* ≤ 0.0001, Pearson’s correlation. MESA, n=1202; BSGS, n=614; GMPWAR, n=656; SATSA, n=1072.

To elucidate the relationship between the expression and methylation level of RTEs, we divided the MESA samples, with matched RTE DNA methylation and expression, into three groups based on the level of expression of RTEs in different classes and families: low (1^st^ quartile), medium (2^nd^ and 3^rd^ quartile), and high (4^th^ quartile) expression groups of monocyte samples from the MESA cohort. The DNA methylation levels of LINEs and LTRs significantly decreased in groups of high LINE and LTR expressions, respectively, but SINEs and the satellite repeats (as negative control) did not display the same pattern (**Fig. 4c**). LINE families, MIR, and LTR families except ERVK displayed lower levels of methylation in higher expression groups, while such pattern was absent in Alu, CR1, and ERVl (**Supplementary Fig. 11**). Although the DNA methylation level of SINEs positively correlated with the methylation levels of LINEs and LTRs, the expression of SINEs was not significantly correlated with the expression levels of other RTE classes (**Fig. 4d**). Overall, we found a distinct pattern for SINEs versus LTRs and LINEs by exploring a relationship between DNA methylation and expression levels of RTEs in monocytes, consistent with our other findings showing different patterns for SINEs versus LTRs and LINEs in PBMCs (see above). This result also suggests a dispensable role of DNA methylation for SINE silencing in monocytes, which is in line with previous studies showing SINEs are not derepressed by deletion of DNA methyltransferases or treatment with DNA demethylating agents and are primarily regulated by histone modifications^34^.

### Elevated level of RTEs in plasma cells of healthy PBMC samples is associated with the high SASP and inflammaging gene signature scores

As described above, we did not identify any association between RTE expression and chronological aging for any human cohorts we analyzed. However, the positive correlation between LINE and LTR expressions with BAR gene signature scores in the PBMC samples of the GARP cohort prompted us to investigate PBMC samples in more detail using PBMC scRNAseq data of 21 samples annotated into 25 cell types by Mogilenko et al.^15^(**online methods** and **Supplementary Table 6**). The scRNA-seq samples were divided based on low and high BAR gene signature scores to compare against the expression of RTE classes for each cell type (**online methods**). Among the 25 annotated cell types, plasma cells consistently demonstrate significantly elevated RTE expression in cells of high SASP expression that is not observed in the other cell types (**Fig. 5a** and **Supplementary Figs. 12-14)**. Increased LINE and SINE expressions were also observed in the plasma cells of the samples with high inflammatory chemokines and inflammaging gene signature scores (**Fig. 5b** and **c**). Although not statistically significant, elevated RTE expression was also seen in the groups with high inflammatory cytokines and senescence but not IFN-I scores (**Fig. 5d-f**). Overall, this finding indicates that plasma cells might play an important role in the process in which reactivation of RTEs regulates aging.

**Figure 5.**
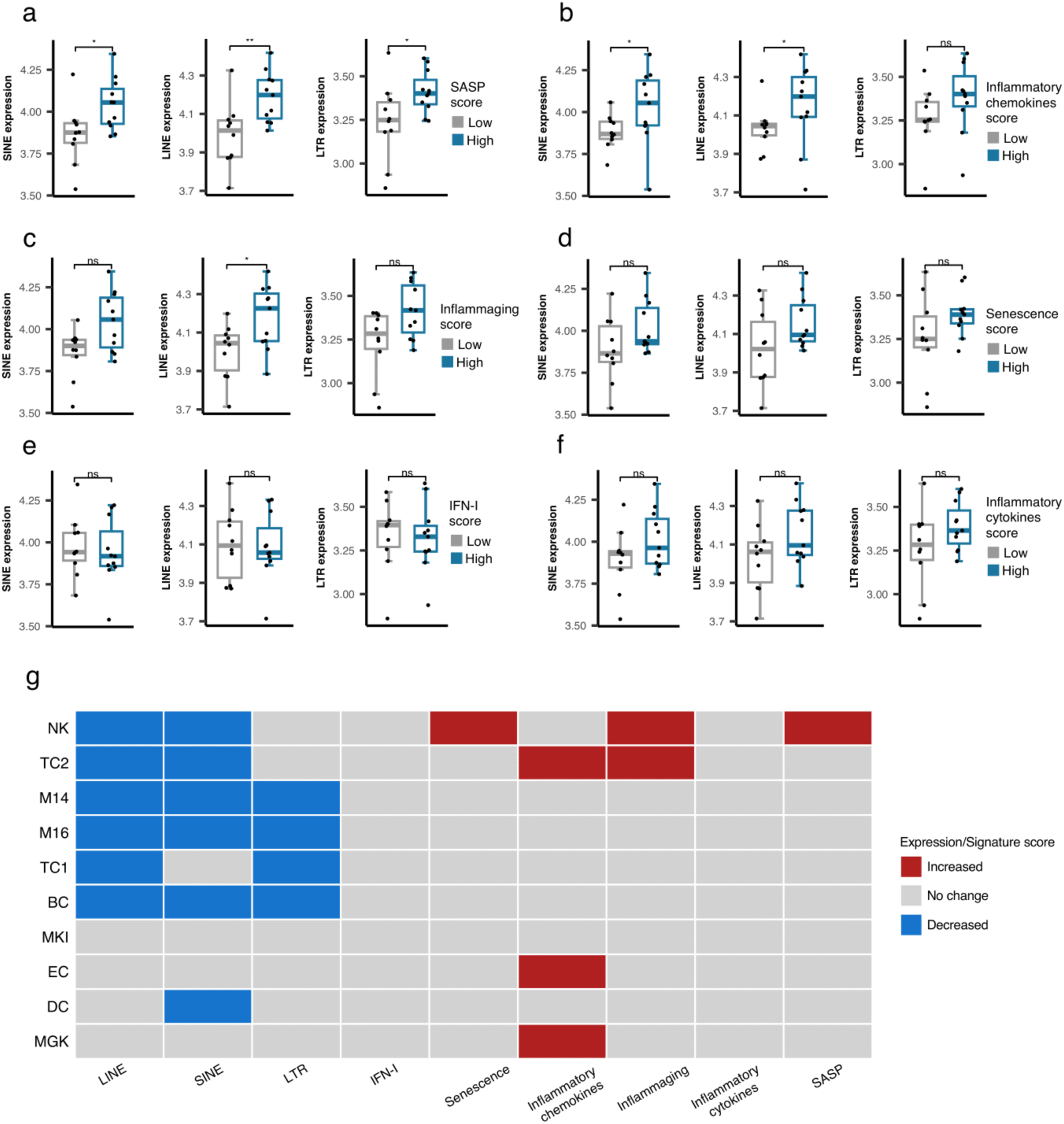
Cell type-specific analysis of RTE expression vs BAR gene signature scores in two PBMC scRNA-seq cohorts. **a-f**, Unique increased expression of RTEs in the Plasma B cells of the samples with high BAR gene signature scores indicates the potential role of plasma B cells in aging. Wilcoxon test. n = 21. **g,** Decreased expressions of RTE classes and increased BAR gene signature scores in multiple annotated cell types obtained from the PBMCs of supercentenarians compared to ordinary elderlies as control. Supercentenarians, n=7; control, n=5, age 50 to 80. Wilcoxon test was applied to identify the significant changes. NK, Natural killer cell; BC, B-cell; TC1, T-cell 1; TC2, T-cell 2; M14, CD14+ monocyte; M16, CD16+ monocyte; EC, Erythrocytes; MKI, MKI67+ proliferating cell; DC, Dendritic cell; MGK, Megakaryocyte.

### Downregulation of RTE expression in supercentenarians versus normal aged cases despite high BAR signature scores in NK and T cells of supercentenarians

Supercentenarians who have reached 110 years of age are an excellent resource for investigating healthy aging. We hypothesized that supercentenarians should express RTEs less than normal-aged people because RTE expression, particularly LINE and LTR expressions, can activate the inflammatory pathways (**Fig. 3**), which would be generally harmful to healthy aging. To understand whether there is any RTE expression change in supercentenarians versus normal-aged people and identify the relationship to BAR gene signatures, we compared the scRNA-seq data of the PBMCs of seven supercentenarians (over 110 years) to five non-centenarian (50 to 80 years) controls^17^. We collected the annotated cells of these twelve cases as a Seurat object for which the cells were annotated into ten different cell types. Compared with the control group, the supercentenarians demonstrated decreased RTE expression in the natural killer (NK) cells, B-cells, T-cells, monocytes, and dendritic cells (**Fig. 5g** and **Supplementary Fig. 15**). Both T cell types showed decreased LINE expression, while the noncytotoxic cluster, TC1, showed slightly decreased LTR expression, and the expanding cytotoxic T cells, TC2, showed decreased SINE expression. On the other hand, the cytotoxic T cells also demonstrated an increased level of inflammatory cytokines and inflammaging gene signature scores. In addition, the NK cells displayed higher senescence and SASP scores in the supercentenarians while demonstrating decreased LINE and SINE expressions (**Fig. 5g** and **Supplementary Figs. 16-17**). Overall, this result reveals inflammatory activity in cytotoxic T cells and increased senescence in the NK cells that is not associated with the activation of RTEs in supercentenarians. By contrast, the expression of RTEs is significantly reduced in most of the immune cell types in supercentenarians.

## Discussion

The role of RTEs in aging has been studied extensively but not in a systematic manner with large, non-cancerous human cohorts across RTE categories before this study. Current understanding of the relationship between RTEs and aging are largely conducted *in vitro* or in model organisms^7,9,35^ that might not apply to human due to intraspecies differences^36^. Moreover, the link of RTEs to aging in non-cancerous human cohorts has not been studied systematically in large cohorts.

We combined bulk and single-cell transcriptomic data to examine the relationship between RTE expression versus chronological and biological aging, demonstrating that chronological aging is not significantly linked with RTE reactivation. In this process, we discovered that LTR and LINE expressions are positively correlated with the BAR gene signature scores. However, in contrast, the SINE expression demonstrates an inverse correlation with BAR scores in the PBMC samples and no significant relationship in the WB. We suspect the absence of a positive correlation between BAR scores and SINE expression might be linked to the same discrepancy observed in the GSVA analysis, in which DNA repair pathways are upregulated in the groups of cases with high SINE and Alu expressions. This result suggests that reactivation of SINE, particularly Alu, might lead to DNA damage, a hallmark of aging. Although SINEs compose up to 13% of the human genome, and Alu elements make up the largest family of human mobile elements, Alu is not studied as extensively as L1 from the LINE class^1^.

Therefore, this novel discovery requires further study to confirm the connection and elucidate the mechanism by which DNA repair might be linked to the derepression of SINE and Alu expressions. In contrast, the groups of cases with high LINE and LTR expressions have upregulation of the pathways related to the inflammatory response, which is in line with the previous findings from human and model organisms^3,35^.

The mechanisms of aging and age-related diseases (ARDs) converge on inflammaging^37^. Previous studies have shown RTE’s pathogenetic role in ARDs including neurodegenerative diseases through elevation of DNA damage and genome instability^38–40^. SINE expression inversely correlated with the BAR signature scores in the PBMCs of osteoarthritis patients, while a positive correlation is observed in monocytes. Moreover, a discrepant upregulation of DNA repair pathways is seen in the osteoarthritis cohorts expressing higher level L2, ERV1, Alu, and MIR. With osteoarthritis being a typical ARD, our analysis provides insight into the relationship between aging and RTEs in an ARD state in comparison with that of non-ARD individuals.

The relationship between DNA methylation and expression of RTEs with age was also investigated in our study. The methylation levels of all three RTE classes decrease with chronological age in our analysis, and the RTE expression is in inverse correlation with the methylation level in LINE and LTR but not in SINE. RTE sequences are repressed via DNA methylation and heterochromatinization^41^. In the aging process, surveillance of transcriptional regulators such as SIRT6 wanes, leading to hypomethylation and heterochromatin reduction^4^. L1 has been identified to be repressed through DNA methylation^42^. However, SINE has been suggested to be predominantly regulated by histone modification, and loss of DNA methylation has little effect on the transcription of SINEs^34^, as reflected in our analysis.

Expression microarray has been successfully employed to study the expression of RTEs43-45. However, the probe design of microarrays does not accommodate specifically to the repeIIve and interspersed nature of RTEs^46^. To circumvent this issue, our analysis was conducted on the class and family levels but not subfamily levels of RTEs to retain enough probes (**Supplementary Fig. 1**) providing sufficient power for calculating the expression scores. We also supported our findings with scRNA-seq, particularly when investigating the association between RTE expression and chronological age, to circumvent the potential constraints posed by using microarray data. However, we are aware of the limitations imposed by using microarray in this study, particularly the low number of RTE probes in the expression microarray data. Although we included one recently published RNA-seq dataset to validate our microarray result, our study can be enriched with the advent of large RNA-seq cohorts for aging studies in the future.

Our exploration of multiple cell types based on a published PBMC scRNA-seq cohort unveiled plasma cell as the only cell type for which the expression of RTEs correlates with BAR signature scores. This matches a recent finding showing a late senescent-like phenotype marked by accumulation of TEs in plasma cells during pre-malignant stages^47^. Lastly, we explored a supercentenarian PBMC scRNA-seq cohort as a model for healthy aging and identified that RTE expression is generally decreased in supercentenarians. However, the BAR signature scores, particularly senescence and SASP scores, increased in the NK cells of supercentenarians compared to the normal-aged group, suggesting that immunosenescence is an important factor in aging.

## Acknowledgment

This work was supported by the Matt Wilson Scholarship from the Faculty of Life Sciences & Medicine, King’s College London. This study was supported with funding from Bristol Myers Squibb (BMS) company during this project. The scRNA-seq raw and processed data of healthy aging populations and the supercentenarians were provided by Maxim N. Artyomov from Washington University School of Medicine in St. Louis and Piero Carninci from RIKEN Center for Integrative Medical Sciences, Yokohama, respectively. We thank Dr D. Mager for critical reading of the manuscript.

## Code availability

Scripts and data used in this study are available on Github (https://github.com/Karimi-Lab/TE_aging_manuscript).

## Methods

### Illumina HT12-v4 probes

The probe lists of Illumina Human HT-12 V3 (29431 probes) and V4 (33963) share 29311 probes in common, which is 99.6% of the list of V3. Therefore, we proceeded with HT-12 V4 probes throughout the analysis to cover all studies that used either V3 or V4. To identify the Illumina probes covering RTE regions, we first selected the Illumina Human HT-12 v4 probes covering intergenic or intronic regions, obtaining a list of 924 unique probes. We then overlapped these probe locations with RepeatMasker^14^ regions to acquire the probes covering RTE regions. In total, we could find 232 unique probes covering RTE regions.

### Generating gene signature scores using *singscore*

For each of the curated gene sets, instead of looking at individual gene expressions, we used *singscore* (version 1.20.0)^31^, a method that scores gene signatures in single samples using rank-based statistics on their gene expression profiles, to calculate the gene set enrichment scores. We first compiled the microarray expression matrix for average expression values from RPM values using *limma*^48^ package in R, then we used the *rankGenes* function from the *singscore* package to rank each gene sample-wise. Eventually, the *multiScore* function was used to calculate signature scores for all six gene sets at once.

### Statistical analyses

All statistical analyses were performed in R (version 4.3.0). Pearson correlation test was used to determine the r and *P* values in correlation analyses. For differential expression analysis, the Wilcoxon test is used to determine the significance. The threshold to determine significance is set at *P* value < 0.05.

### Gene Set Variation Analysis (GSVA)

GSVA was performed using the *GSVA* package (version 1.48.3)^32^ in R to compare the groups of low and high RTE expression groups. The gene lists were retrieved from the Molecular Signatures Database (MSigDB)^33^ by searching for “inflammatory” and “dna_repair” as keywords. We further removed pathways that ended with “_DN” (as for downregulated) or “_UP” (as for upregulated) to reduce repeat and confusion. Three repetitive gene sets were removed from our analysis (GOBP_POSITIVE_REGULATION_OF_ACUTE_INFLAMMATORY_RESPONSE, GOBP_POSITIVE_REGULATION_OF_CYTOKINE_PRODUCTION_INVOLVED_IN_INFLA MMATORY_RESPONSE, GOBP_POSITIVE_REGULATION_OF_ACUTE_INFLAMMATORY_RESPONSE_TO_ANTIGENIC_STIMULUS). The filtering process generated 50 gene sets for inflammatory response and 21 gene sets for DNA repair. To detect any alterations in gene expression, the log fold change threshold was kept low at |logFC| > 0.1, and a significance threshold of *P* < 0.05 was set.

#### ScRNA-seq analysis workflow for RTEs

Two single-cell transcriptomic datasets of healthy human cohorts were adopted in this study (**Supplementary Table 6**). To obtain the RTE expression in single-cell sequencing (scRNA-seq) data, the scRNA-seq bam files were processed through scTE^49^ pipeline. More specifically, for each cell in each sample, the read counts for different classes and families of RTEs were generated using the scTE method. Subsequently, in each sample, we calculated the cumulative read counts for each RTE class per annotated cell type by adding the number of reads belonging to all the cells of each cell type. This resulted in pseudo-bulk read counts of RTE classes for different cell types in each sample.

In parallel, we also calculated the pseudo-bulk read counts for all genes per cell type in each sample using the genic read counts embedded in Seurat^50^ objects collected for the two scRNA-seq datasets that we employed in our study (**Supplementary Table 6**). Concatenating pseudo-bulk RTE and gene read counts, we performed Reads Per Million (RPM) normalization per cell type per sample. For each cell type, the denominator for RPM normalization was the cumulative counts of genes and RTE classes for that cell type.

To calculate the age-associated gene signature scores, we converted the scRNA-seq read counts to bulk RNA-seq read counts by summing up the number of reads of all cells of the same type for each gene. This resulted in a bulk RNA-seq gene count matrix from which we calculated each sample’s RPM values per gene. Next, the RPM matrix was provided to *singscore* to calculate gene signature scores. Lastly, we divided the samples into high vs low gene signature scores by using the median as the cut-off.

#### Bulk RNA-seq analysis for RTEs

The raw RNA-seq reads were mapped to RepeatMasker to extract the reads covering the RTE regions, and then aggregated and normalised for each class and family of RTEs to generate reads per million (RPM) scores.

---

**Supplementary Figure 1.**
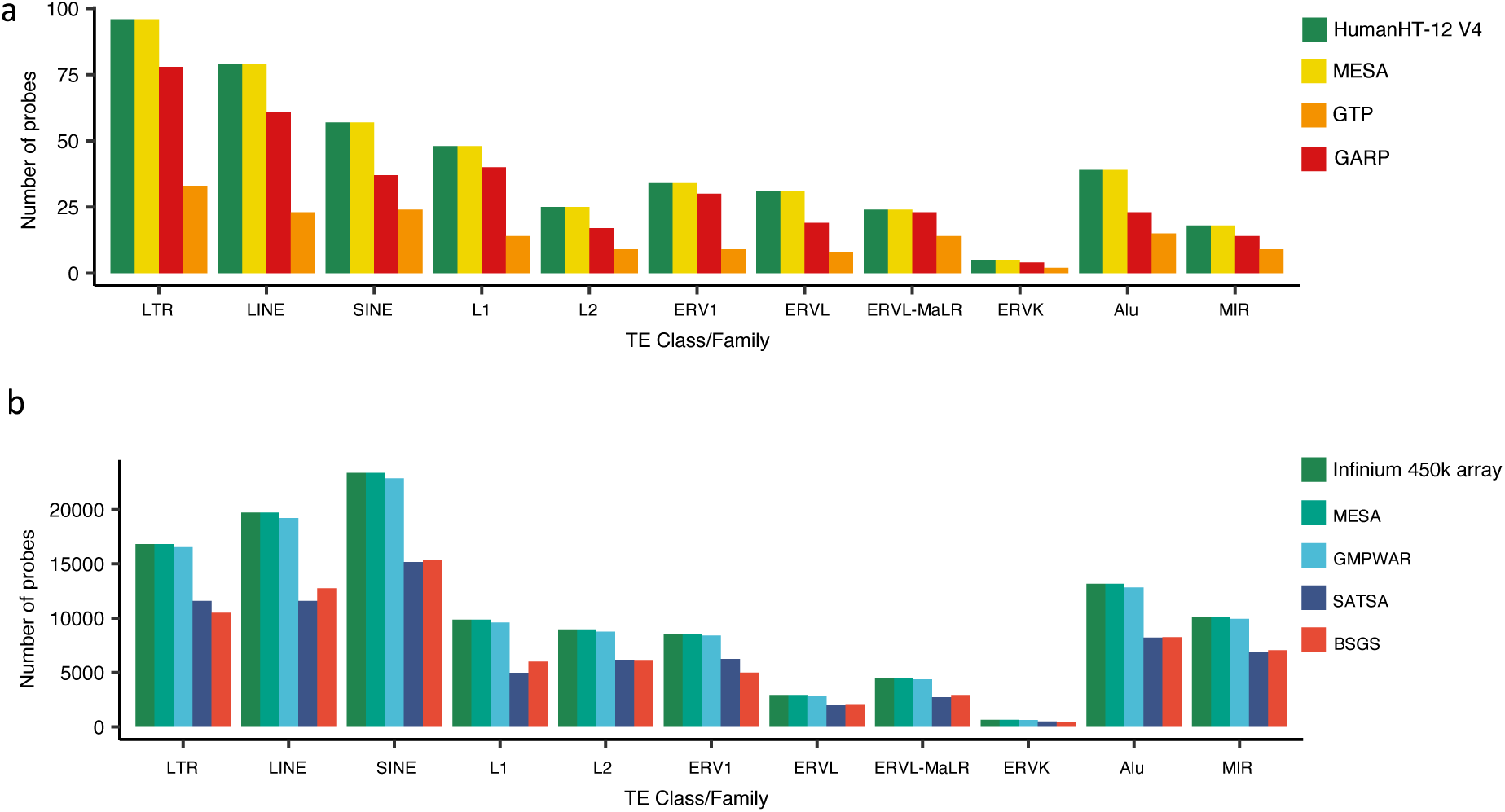
Number of RTE-covering probes. **a**, The number of microarray probes covering RTE regions in MESA, GTP, and GARP compared to the total number of RTE probes available in Illumina HumanHT-12 V4. **b**, The number of microarray probes covering RTE regions in MESA, GMPWAR, SATSA, BSGS compared to the total number of RTE probes available in Illumina Infinium 450k array.

**Supplementary Figure 2.**
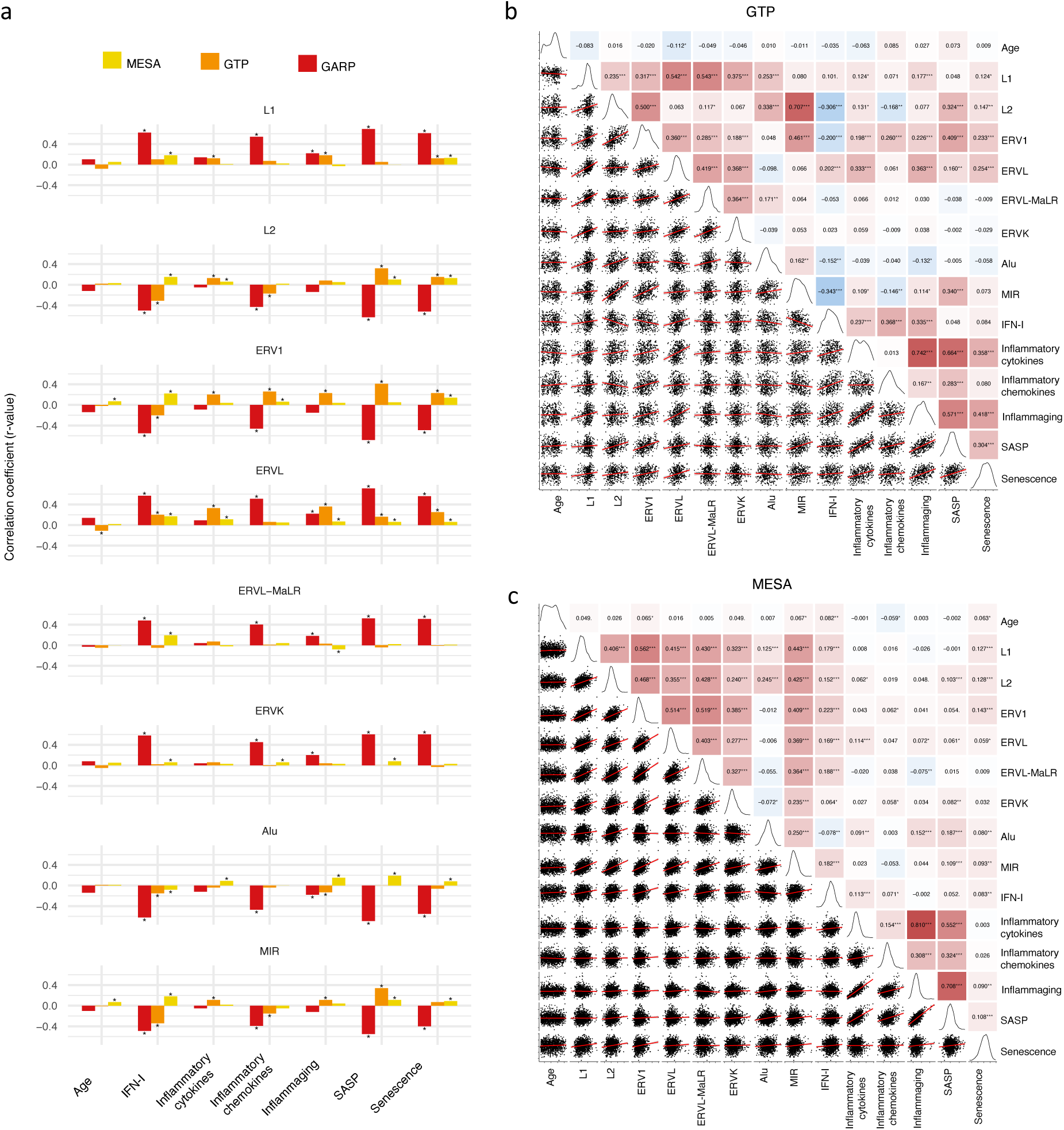
Correlation analysis between BAR gene signatures and RTE family expressions in human cohorts. **a**, Weak correlation between a few RTE families and chronological age and strong positive correlation between L1, ERVL, ERVK, and MaLR with BAR signature scores in PBMC samples (GARP cohort). **b, c**, Correlation matrix depicting all pair-wise combinations to identify the correlation between chronological age, RTE family expressions, and six age-associated gene expressions in monocytes (MESA) and the WB (GTP). * *P* ≤ 0.05, ** *P* ≤ 0.01, *** *P* ≤ 0.001, **** *P* ≤ 0.0001, Pearson’s correlation.

**Supplementary Figure 3.**
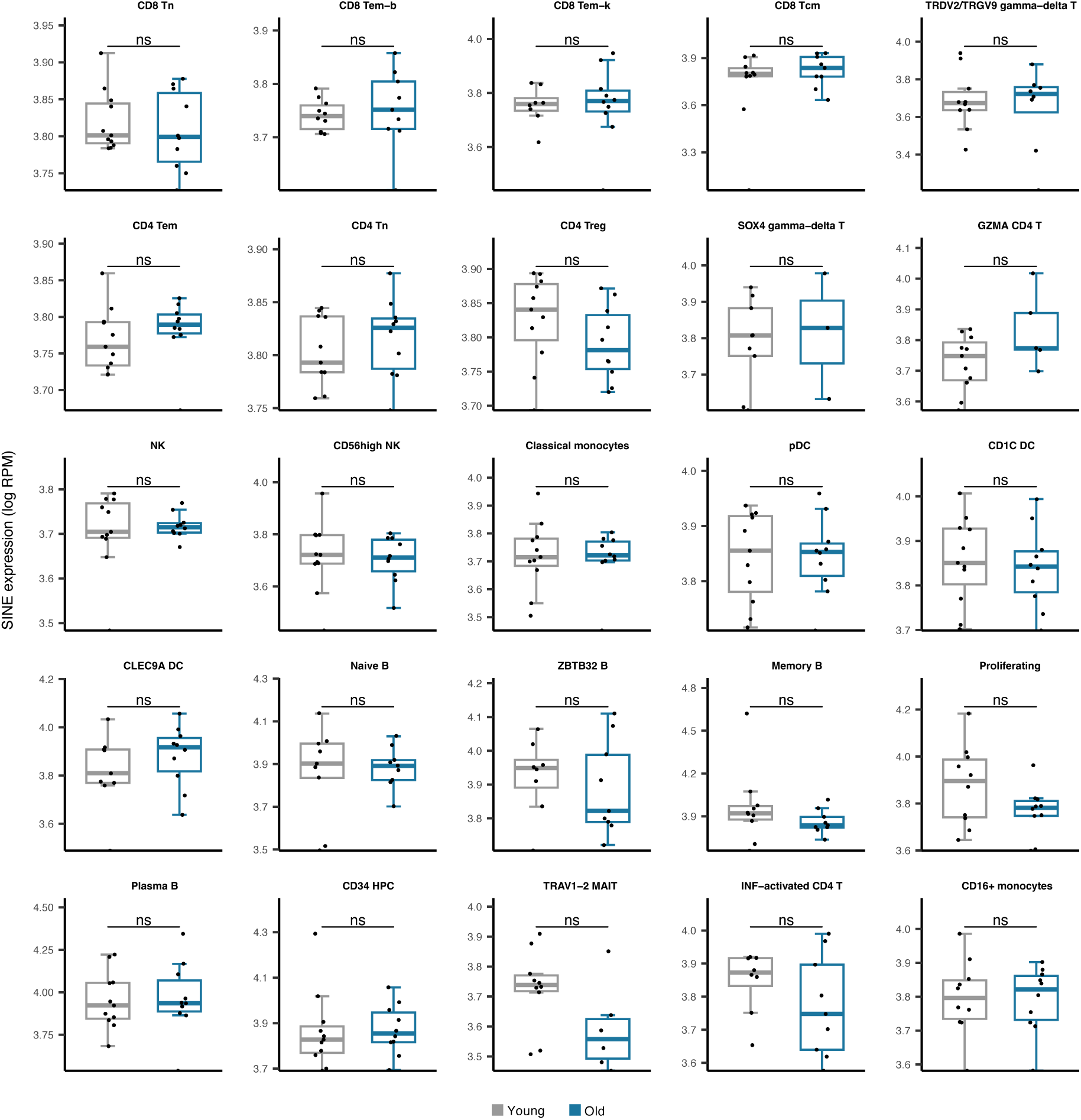
Comparing SINE expression in 25 cell types obtained from young versus old PBMC human samples. No significant difference was observed in any cell type. The young group comprises healthy male donors aged 25 to 29, n=11; the old groups are healthy male donors aged 62 to 70, n=10; Wilcoxon test; ns: insignificant.

**Supplementary Figure 4.**
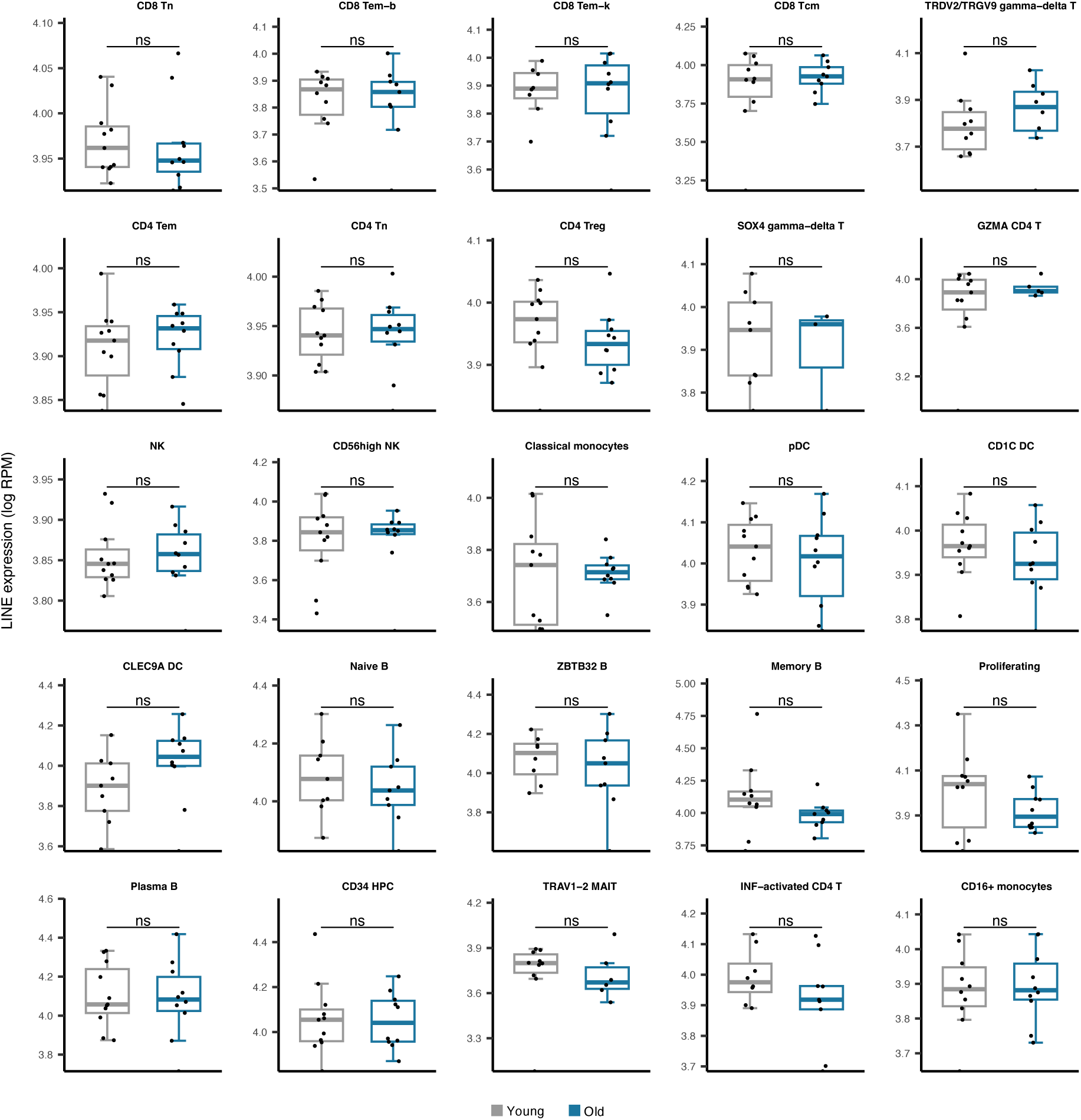
Comparing LINE expression in 25 cell types obtained from young versus old PBMC human samples. No significant difference was observed in any cell type. The young group comprises healthy male donors aged 25 to 29, n=11; the old groups are healthy male donors aged 62 to 70, n=10—Wilcoxon test; ns: insignificant.

**Supplementary Figure 5.**
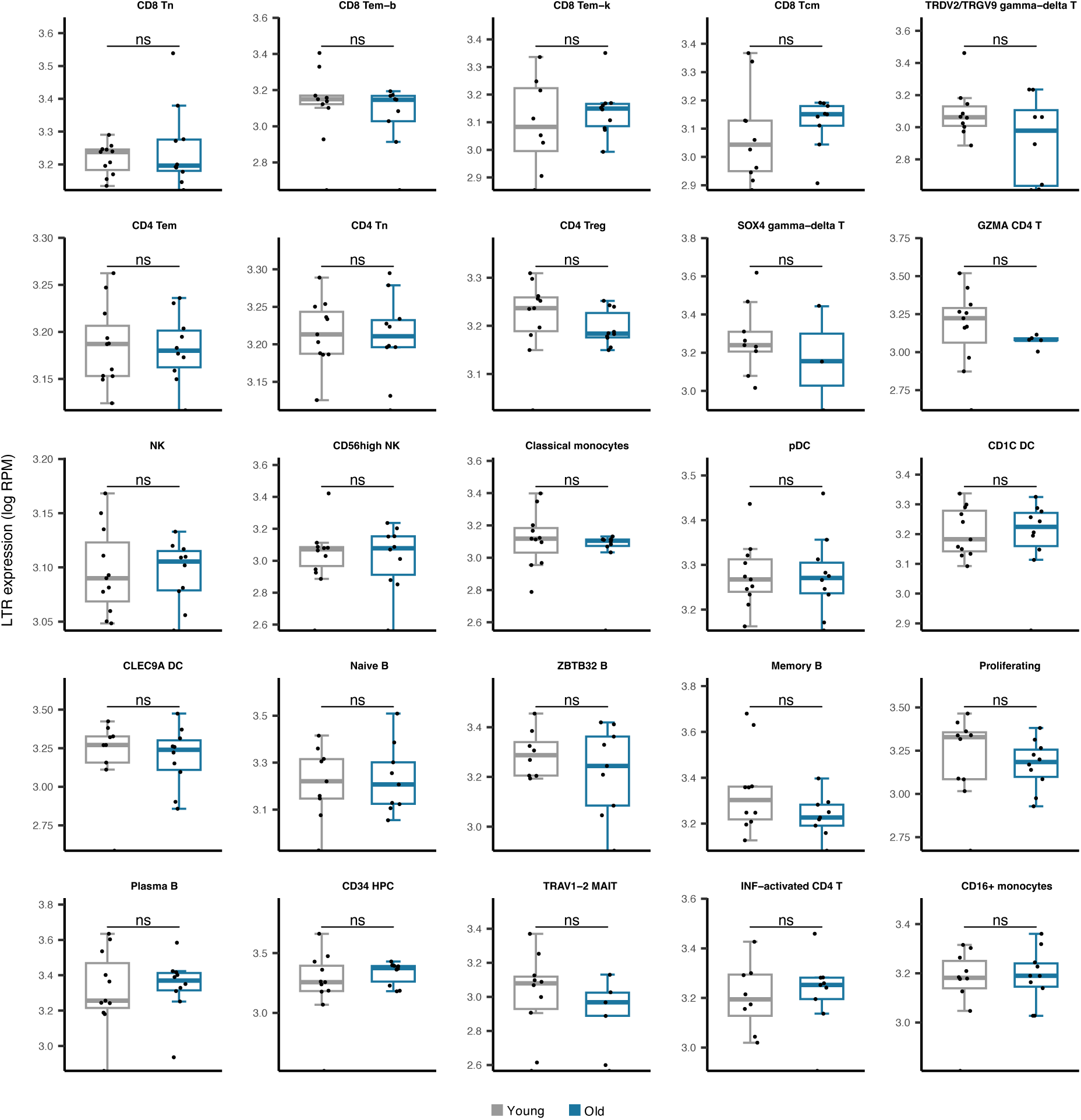
Comparing LTR expressions in 25 cell types obtained from young versus old PBMC human samples. No significant difference was observed in any cell type. The young group comprises healthy male donors aged 25 to 29, n=11; the old groups are healthy male donors aged 62 to 70, n=10—Wilcoxon test; ns: insignificant.

**Supplementary Figure 6.**
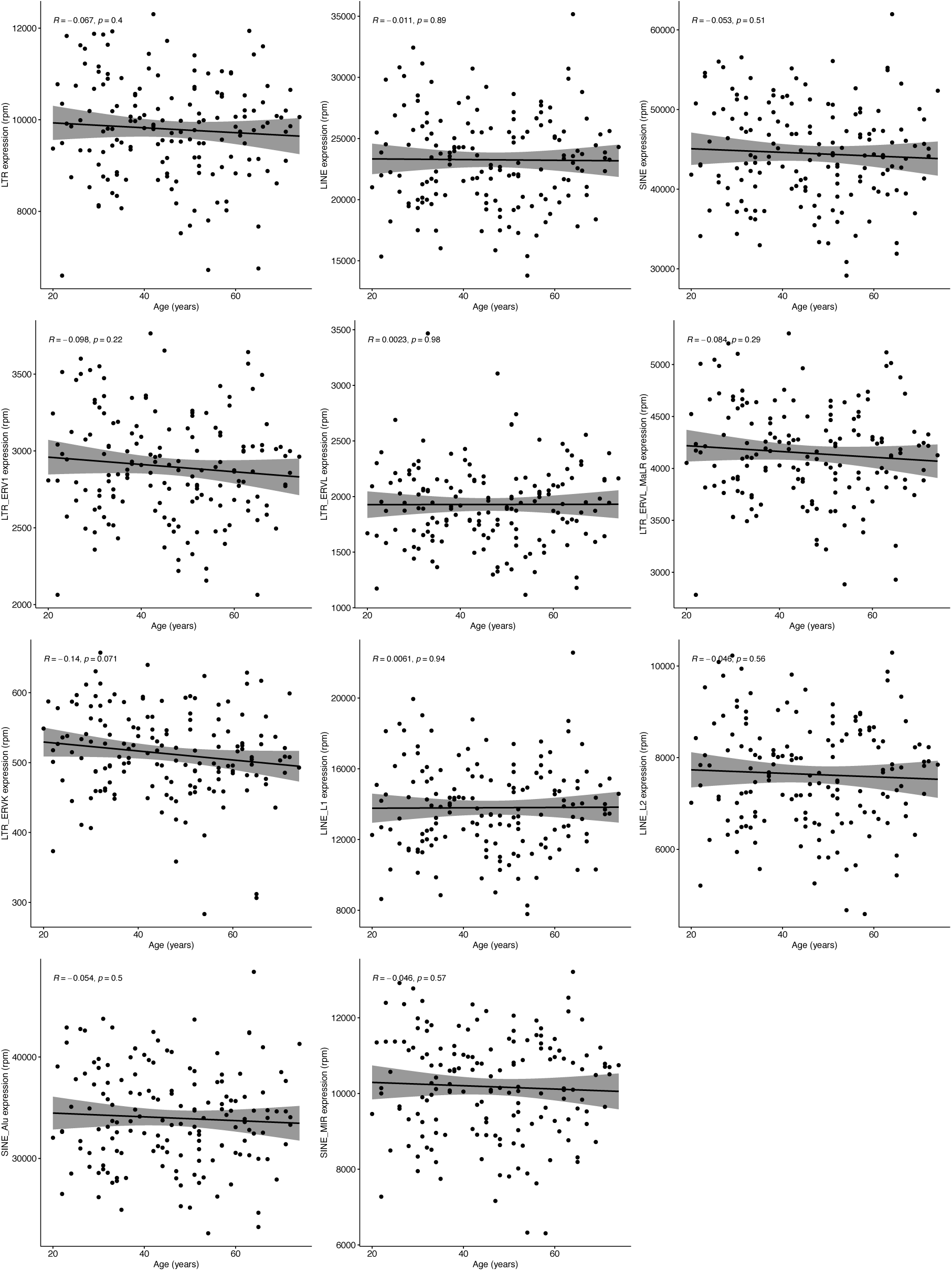
Correlation analysis of RTE expression and chronological age in RNA-seq data of healthy human PBMC samples. No correlation was observed between chronological age and expression of RTE classes or families. Blood samples were acquired from healthy donors of 117 males and 42 females aged 20 to 74 years. n=159—Pearson’s correlation.

**Supplementary Figure 7.**
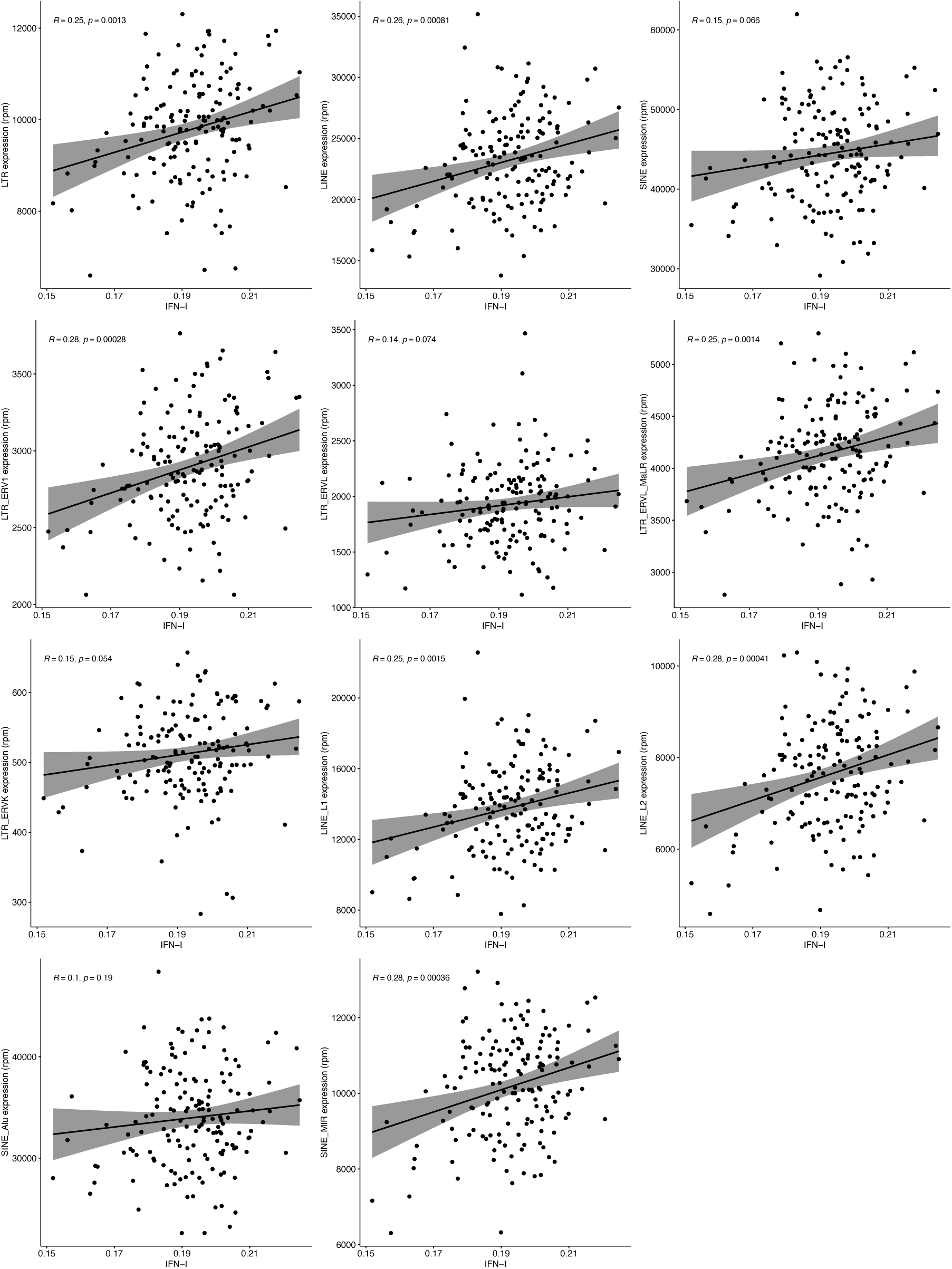
Correlation analysis of RTE expression and IFN-I score in RNA-seq data of healthy human PBMC samples. Despite having a significant correlation between the IFN-I signature score and expression of LINE and LTR classes and most of their families (P-val < 0.01), such correlation did not exist between the IFN-I score and SINE class and Alu family. Blood samples were acquired from healthy donors of 117 males and 42 females aged 20 to 74 years. n=159—Pearson’s correlation.

**Supplementary Figure 8.**
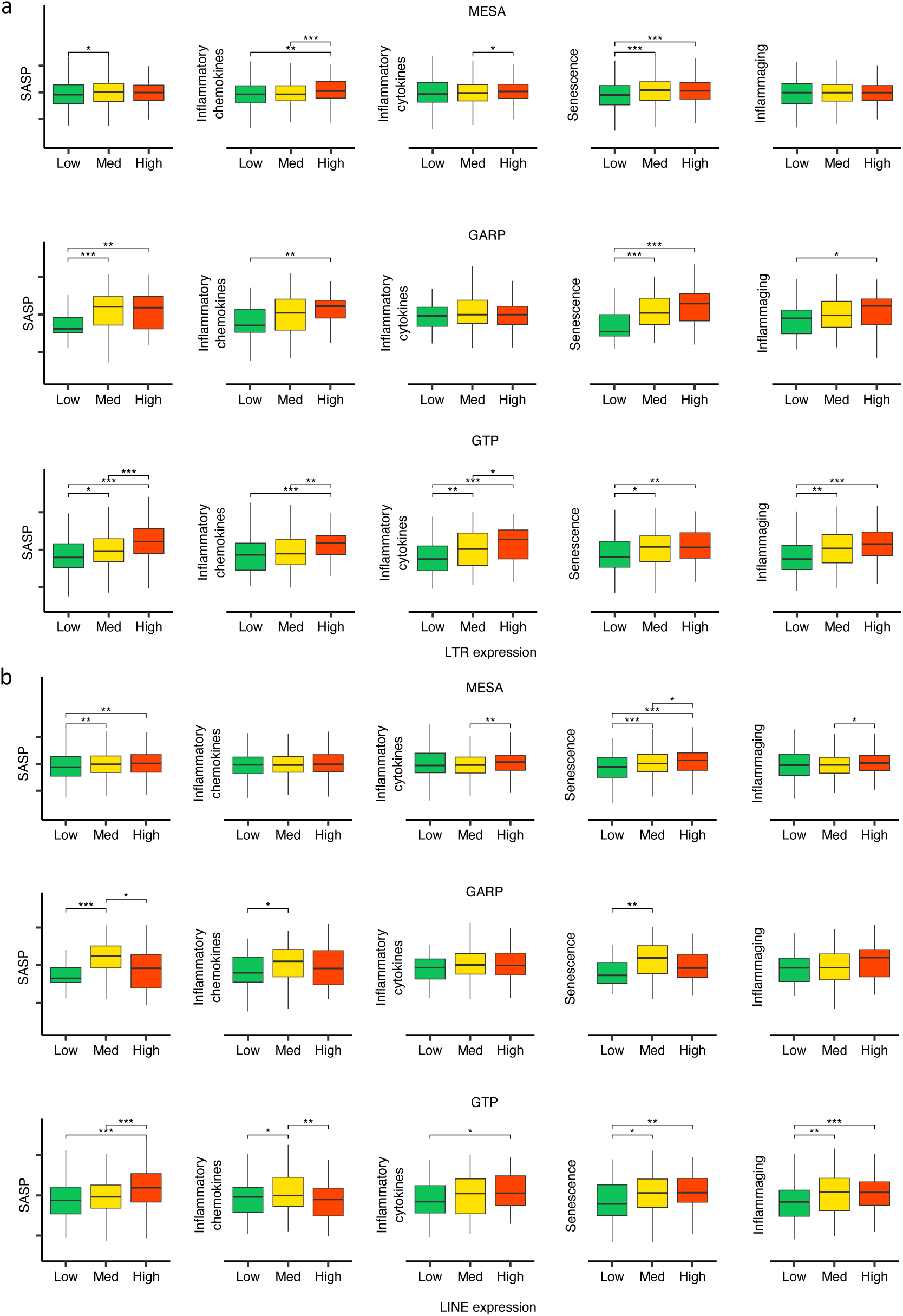
Increasing trend of BAR gene signature scores in high vs low LTR and LINE expression groups in the three human cohorts. **a,b** The samples in each cohort were divided into (low (1^st^ quartile), medium (2^nd^ and 3^rd^ quartile), and high (4^th^ quartile) LTR and LINE expression groups, respectively. * *P* ≤ 0.05, ** *P* ≤ 0.01, *** *P* ≤ 0.001, **** *P* ≤ 0.0001, Wilcoxon test.

**Supplementary Figure 9.**
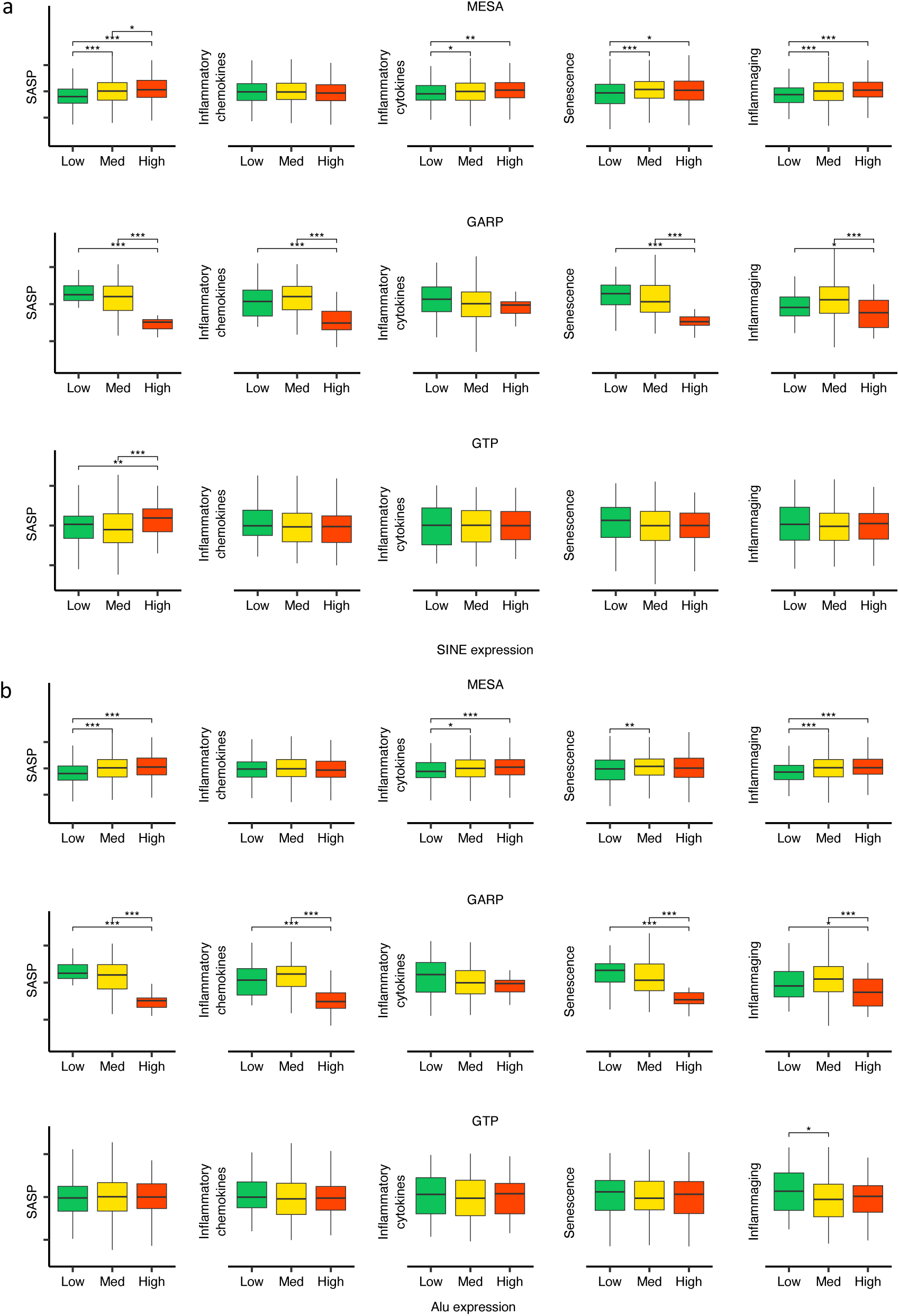
Different patterns of BAR signature scores in MESA vs GARP vs GTP for high vs low SINE and Alu expression groups in the three human cohorts. **a, b**, The samples in each cohort were divided into low (1^st^ quartile), medium (2^nd^ and 3^rd^ quartile), and high (4^th^ quartile) SINE and Alu expression groups, respectively. * *P* ≤ 0.05, ** *P* ≤ 0.01, *** *P* ≤ 0.001, **** *P* ≤ 0.0001, Wilcoxon test.

**Supplementary Figure 10.**
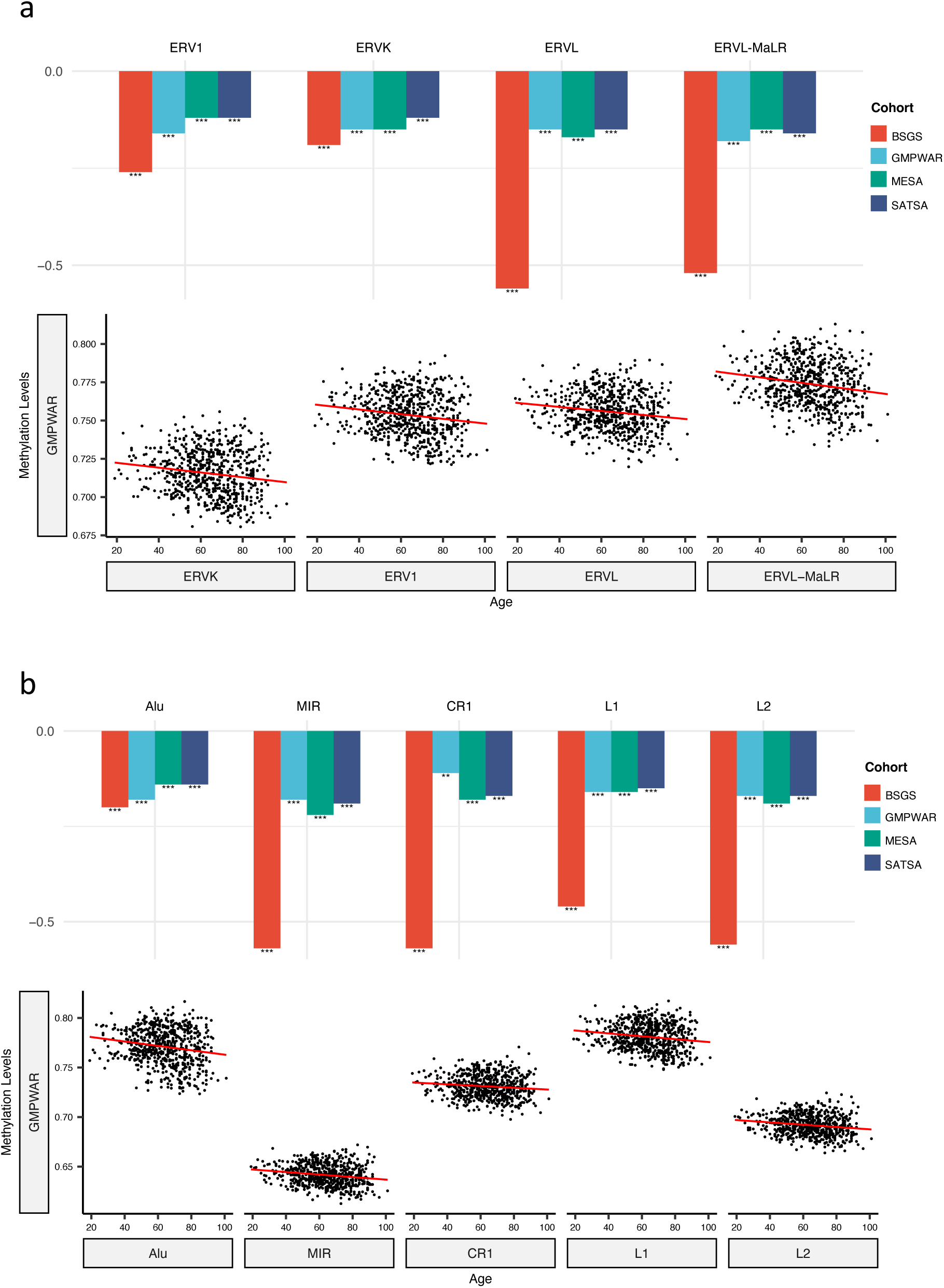
Correlation of methylation levels of RTE families with chronological age in monocyte (MESA) and WB (BSGS, SATSA, and GMPWAR) samples. **a, b**, Methylation levels of LTR and LINE/SINE families negatively correlate with chronological age in monocytes (MESA) and the WB (BSGS, SATSA, and GMPWAR). *** *P* ≤ 0.001, Wilcoxon test.

**Supplementary Figure 11.**
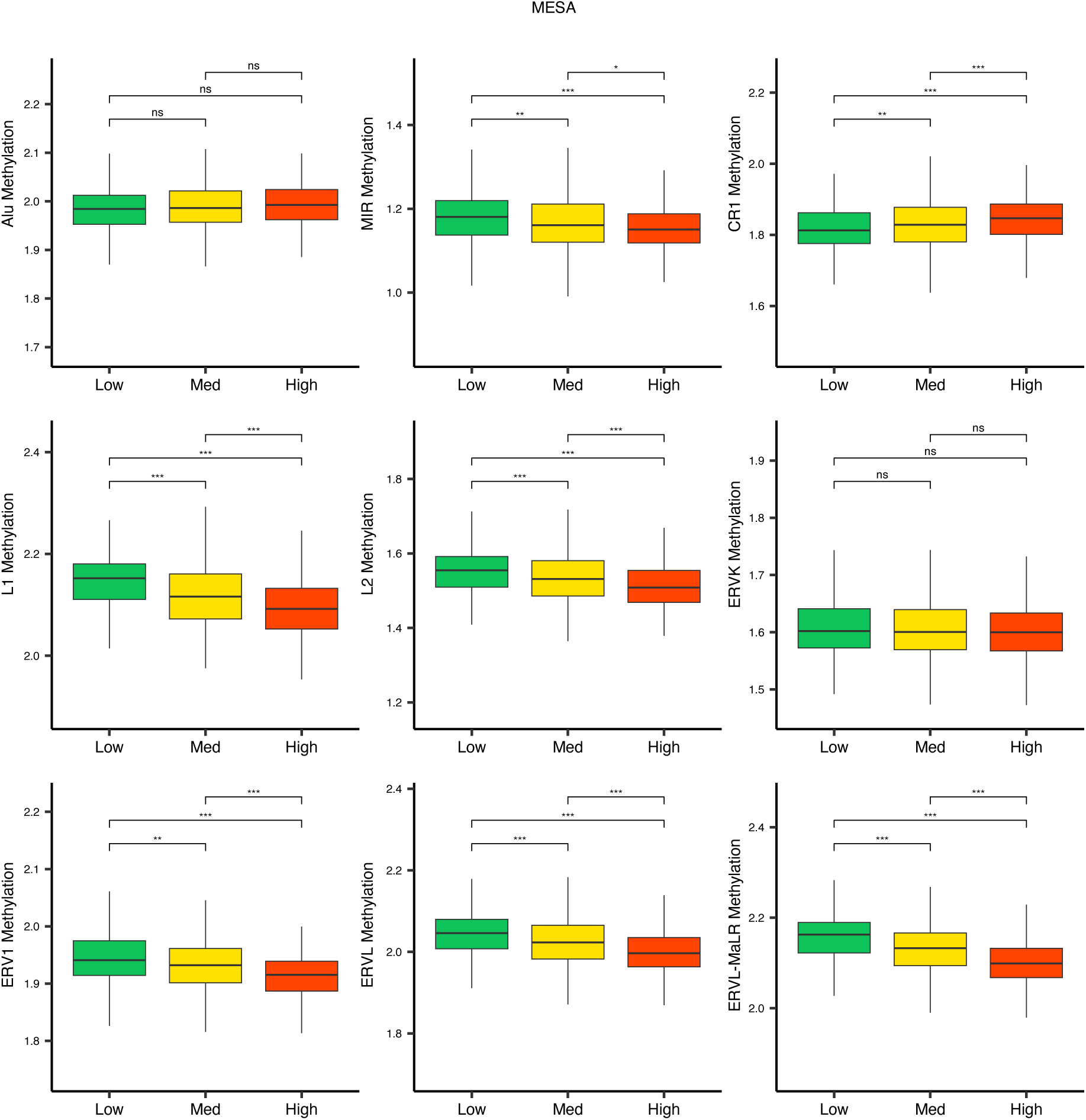
Methylation levels versus low (1^st^ quartile), medium (2^nd^ and 3^rd^ quartile), and high (4^th^ quartile) expressions of RTE families in monocytes. While LINE families, MIR, and LTR families except ERVK show lower levels of methylation in higher expression groups, this pattern is not seen in Alu, CR1, and ERVl. * *P* ≤ 0.05, ** *P* ≤ 0.01, *** *P* ≤ 0.001, **** *P* ≤ 0.0001, ns: not significant, Wilcoxon test.

**Supplementary Figure 12.**
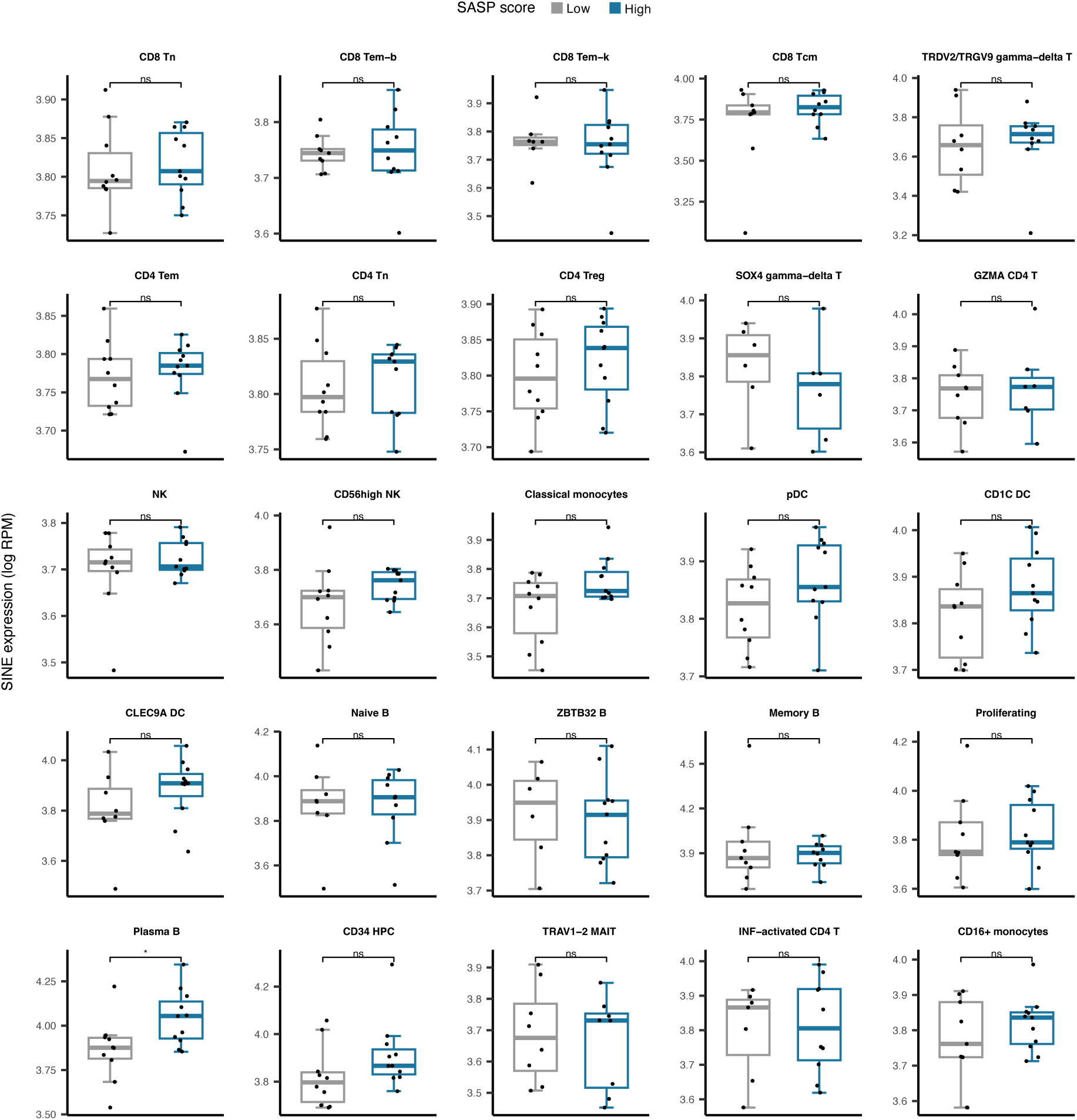
Cell type-specific SINE expression versus low and high SASP score groups among 25 cell types in PBMC. Plasma B cells demonstrate significantly elevated SINE expression in samples with high SASP scores. * *P* ≤ 0.05, ns: not significant, Pearson’s correlation.

**Supplementary Figure 13.**
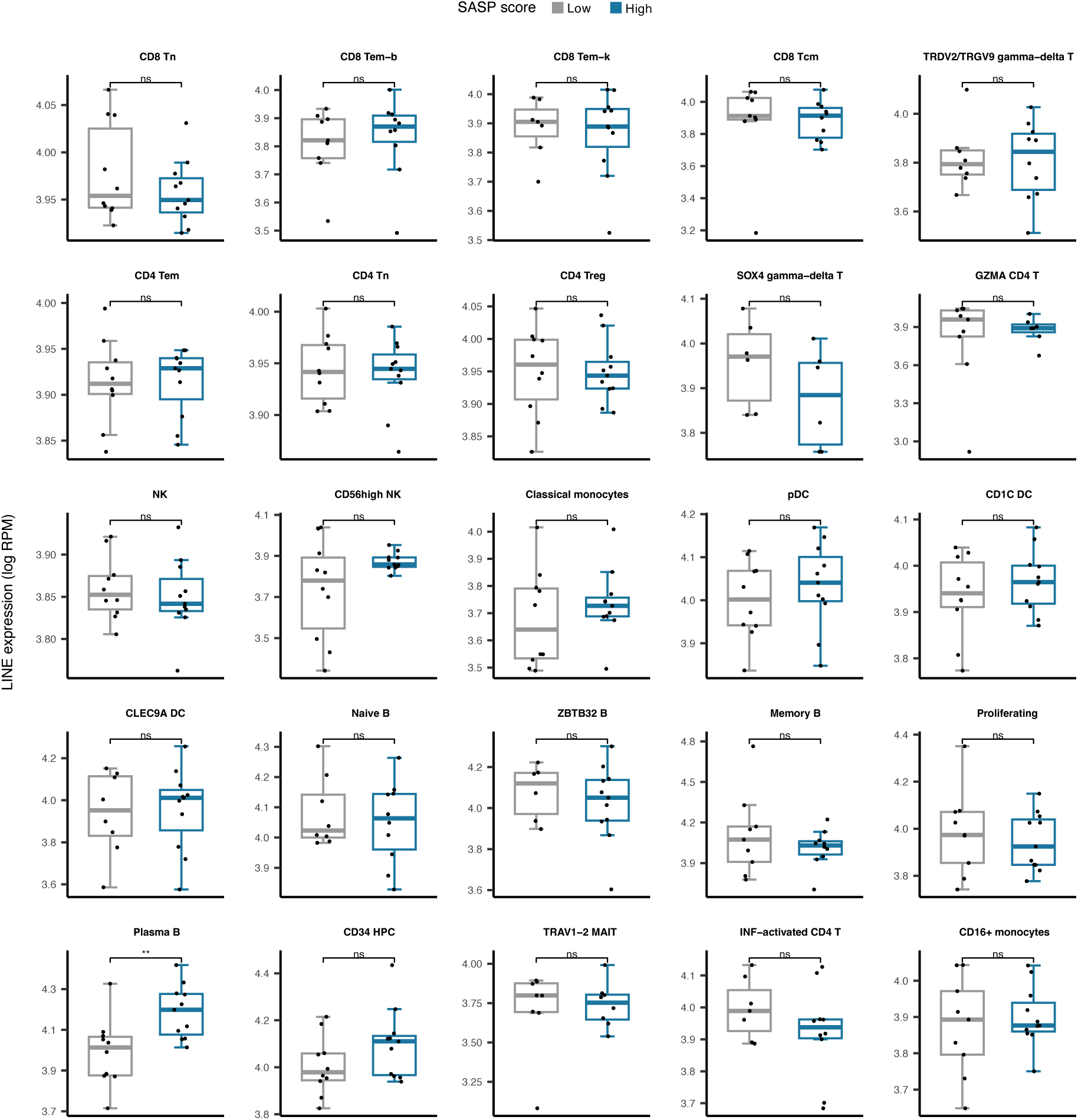
Cell type-specific LINE expression versus low and high SASP score groups among 25 cell types in PBMC. Plasma B cells demonstrate significantly elevated LINE expression in samples with high SASP scores. * *P* ≤ 0.05, ns: not significant, Pearson’s correlation.

**Supplementary Figure 14.**
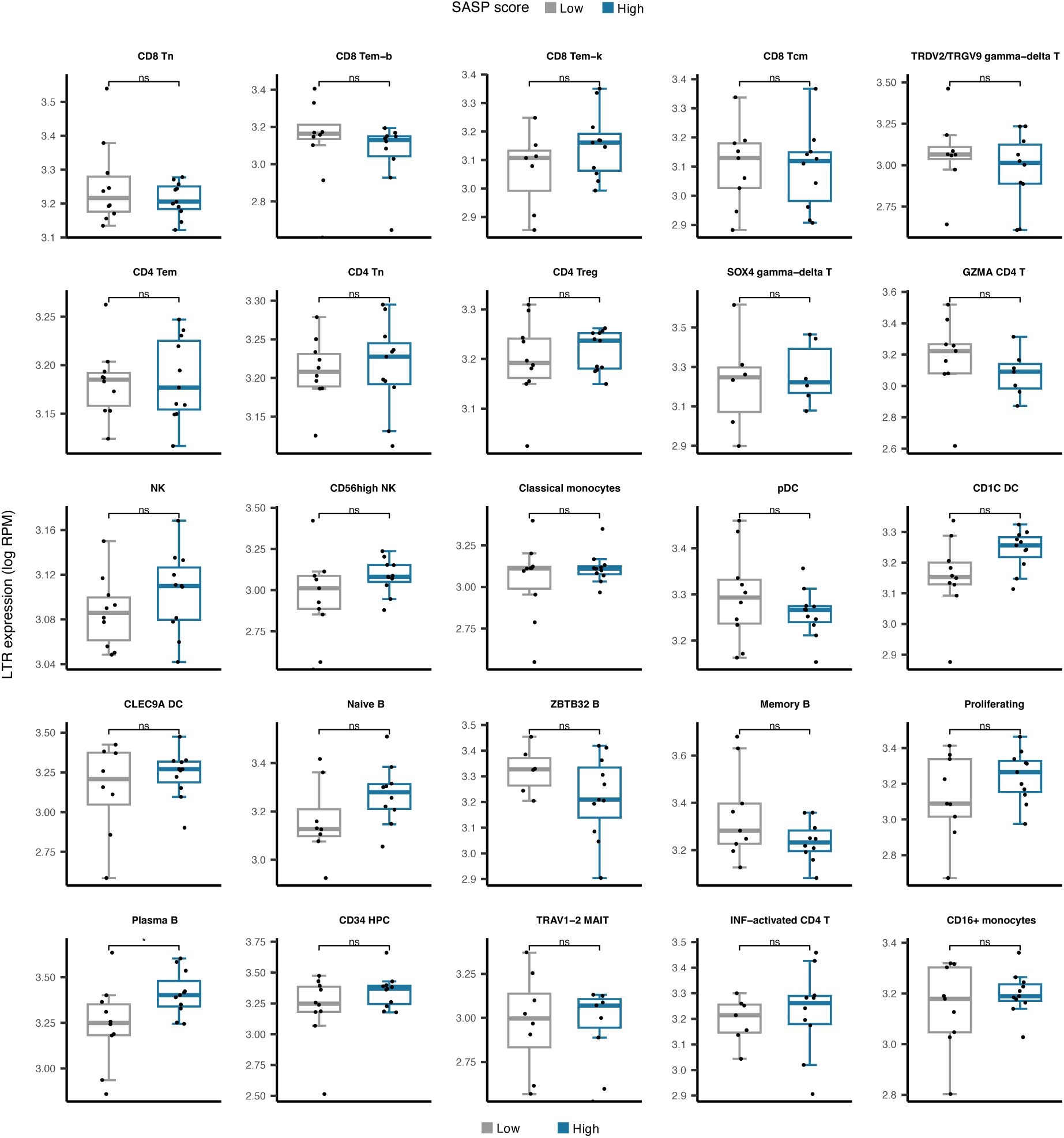
Cell type-specific LTR expression versus low and high SASP score groups among 25 cell types in PBMC. Plasma B cells demonstrate significantly elevated LTR expression in samples with high SASP scores. * *P* ≤ 0.05, ns: not significant, Pearson’s correlation.

**Supplementary Figure 15.**
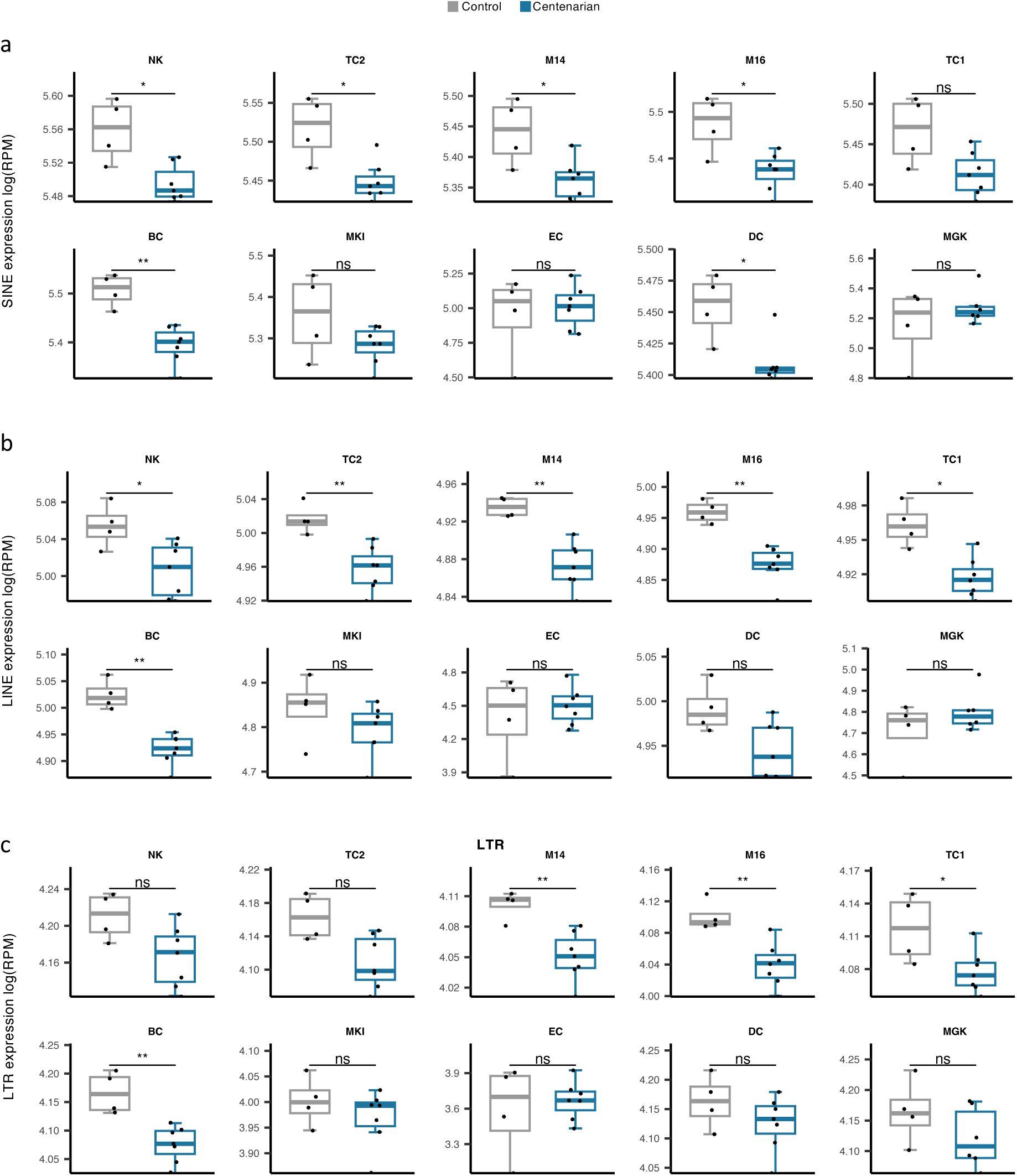
Comparison of cell type-specific RTE expression in supercentenarians versus normal aged cases. * *P* ≤ 0.05, ** *P* ≤ 0.01, *** *P* ≤ 0.001, **** *P* ≤ 0.0001, ns: not significant, Wilcoxon test.

**Supplementary Figure 16.**
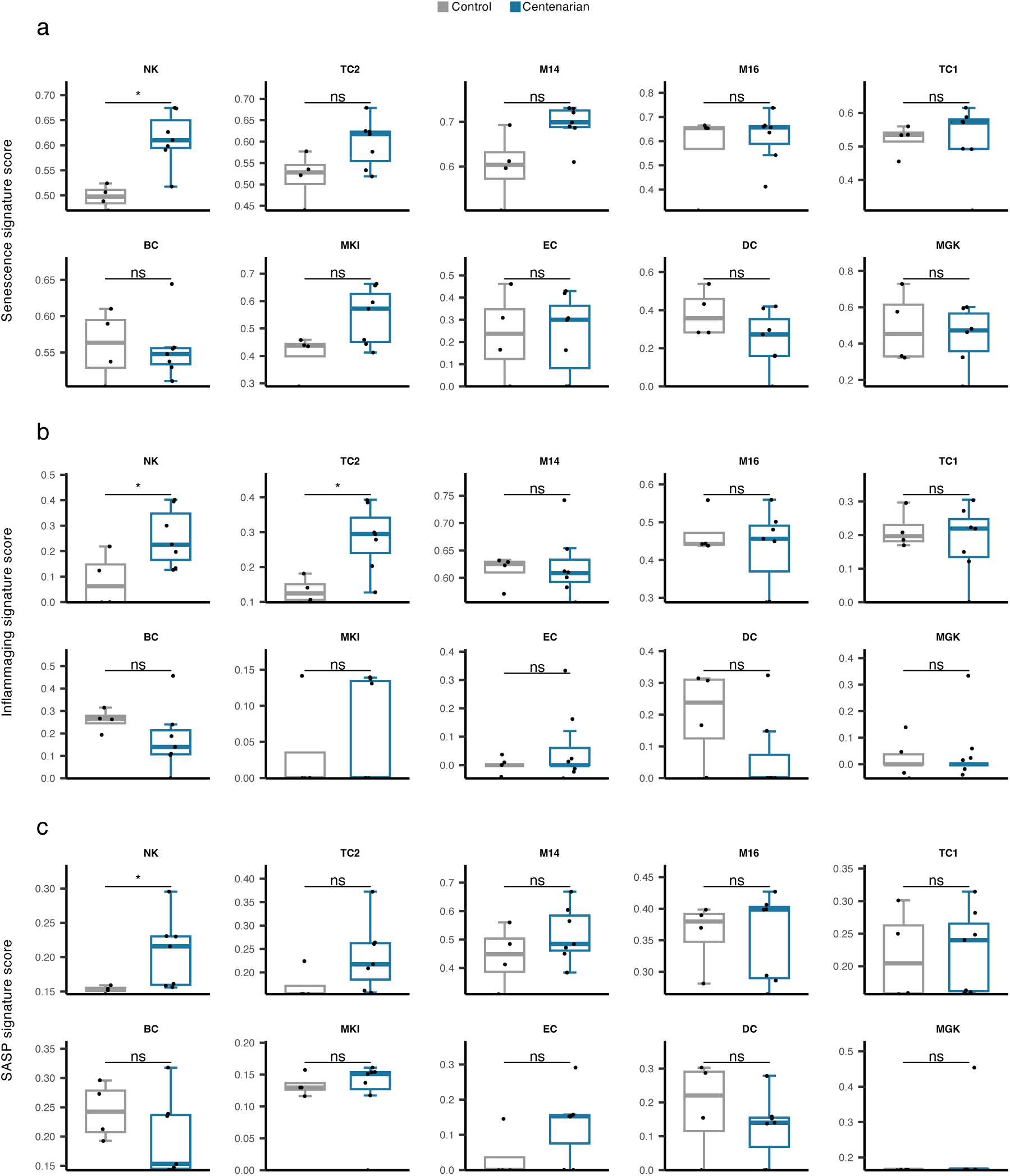
Comparison of cell type-specific Senescence, inflammaging, and SASP gene signature scores in supercentenarians versus normal aged cases. * ***P*** ≤ 0.05, ** *P* ≤ 0.01, *** *P* ≤ 0.001, **** *P* ≤ 0.0001, ns: not significant, Wilcoxon test.

**Supplementary Fig. 17.**
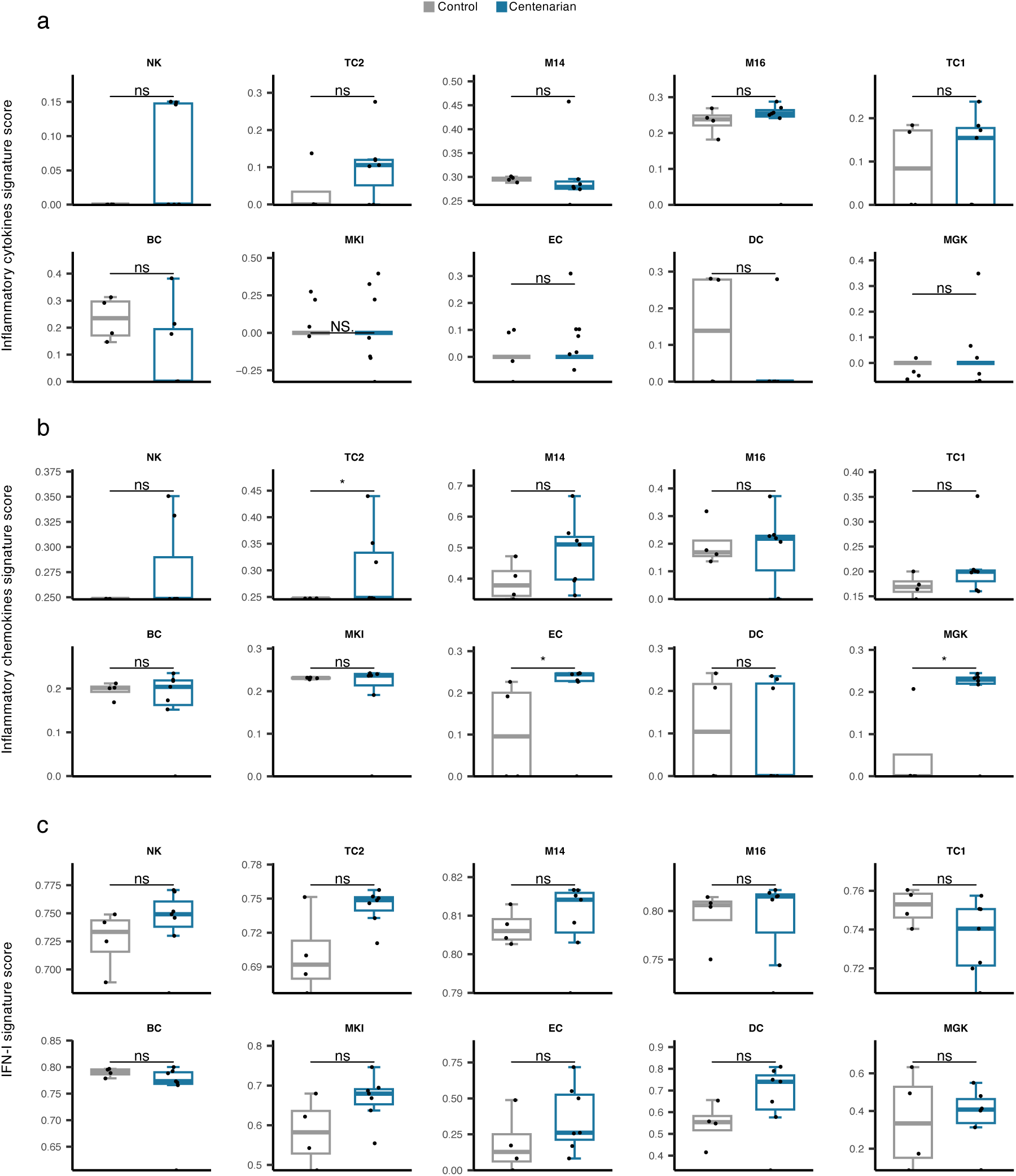
Comparison of cell type-specific Inflammatory cytokines, Inflammatory chemokines, and IFN-I gene signature scores in supercentenarians versus normal aged cases. * *P* ≤ 0.05, ** *P* ≤ 0.01, *** *P* ≤ 0.001, **** *P* ≤ 0.0001, ns: not significant, Wilcoxon test.

## Supplementary Notes

### Descriptions of the study populations included in the analysis

#### Grady Trauma Project (GTP)

The research concentrated on African American individuals of low socioeconomic backgrounds who had undergone trauma, primarily recruited from Atlanta’s Grady Memorial Hospital between 2005 and 2007 (Charles F. Gillespie et al., 2009). Exclusion criteria comprised mental retardation, active psychosis, or inability to provide informed consent. With over 7,000 participants, mostly urban African Americans with low socioeconomic status, the GTP specifically targeted a demographic with a history of significant trauma. The study involved WB collection between 8 to 9 a.m. following overnight fasting. Whole genome expression profiles were generated for 398 subjects at the Max-Planck Institute using Illumina HT-12 v3.0 or v4.0 arrays, focusing on 359 subjects of self-reported African American ancestry aged between 16-78 years. The expression data is available under GSE58137 at the GEO public repository. The Institutional Review Boards of Emory University School of Medicine and Grady Memorial Hospital granted approval for all procedures in this study. Funding primarily came from the National Institutes of Mental Health (MH071537).

#### Genetics, Arthrosis and Progression study (GARP)

The GARP study focused on Dutch Caucasian siblings diagnosed with symptomatic osteoarthritis across multiple joint sites. It comprised 191 sibling pairs (n=382 individuals) of white Dutch ancestry, aged between 40 and 70, diagnosed with primary symptomatic osteoarthritis in multiple joints including hand, spine, knee, or hip (N Riyazi et al., 2005). Whole genome expression profiles were analyzed from 108 participants (68 unrelated families) of the GARP study and 26 age-matched healthy controls. The blood of participants was collected, and mononuclear blood cells were separated prior to RNA isolation. Sample libraries were hybridized onto the microarrays (Illumina Human HT-12_v3_BeadChip’s; Illumina). For the methylation analysis, genomic DNA was extracted, and the DNA methylation was assayed at over 450,000 sites on the Illumina Infinium HumanMethylation 450K BeadChips. The data is available at GEO public repository under the accession GSE48556.

#### Multi-Ethnic Study of Atherosclerosis (MESA)

MESA is a study of subclinical cardiovascular disease and the factors that forecast its progression to clinically evident forms initiated in 2000. This comprehensive investigation involves 6,814 asymptomatic individuals aged 45-84, presenting a diverse demographic: 38% Caucasian, 28% African American, 22 percent Hispanic, and 12 percent Asian (primarily of Chinese descent). Enrolling 6,500 individuals evenly distributed across genders, the study targets those without clinical cardiovascular disease at the outset, representing four racial/ethnic groups from six US communities (Forsyth County, North Carolina; St. Paul, Minnesota; Chicago, Illinois; New York, New York; Baltimore, Maryland; Los Angeles County, California). The study protocol has been approved by the institutional review boards of the six field centers: Wake Forest University, Columbia University, Johns Hopkins University, University of Minnesota, Northwestern University, and University of California - Los Angeles. Detailed information about the design of MESA can be found at https://internal.mesa-nhlbi.org, and the data is accessible under GSE56045 at GEO.

Blood samples were gathered to isolate peripheral blood mononuclear cells, from which monocytes were separated using anti-CD14 monoclonal antibody-coated magnetic beads. Flow cytometry analysis consistently demonstrated monocyte samples of over 90% purity across 18 specimens. The DNA and RNA were extracted simultaneously. RNA with integrity scores above 9.0 was chosen for global expression microarrays. Genome-wide expression analysis employed the Illumina HumanHT-12 v4 Expression BeadChip and Illumina Bead Array Reader, following the Illumina expression protocol. Additionally, the Illumina HumanMethylation450 BeadChip and HiScan reader were utilized for epigenome-wide methylation analysis, assigning individual samples to the BeadChips and chip positions using the same sampling scheme as for the expression BeadChips. Details for sample preparation, data processing and quality control were processed as demonstrated in the design paper (Diane Bild et al., 2002). The expression and methylation data are available at GEO public repository under the accession GSE56045 and GSE56046, respectively.

#### Swedish Adoption/Twin Study of Aging (SATSA)

SATSA is a comprehensive exploration of aging variations using twins raised in different environments influenced by genetic and environmental factors. The study draws its data from the Swedish Twin Registry, a population-based national register encompassing twins born from 1886 to 2000. Additionally, the comprehensive assessment across biological, psychological, and social domains allows for the examination of patterns, predicting the onset of age-related health issues. Started in 1984, the study employed rigorous questionnaires every three years until 2010, engaging over 2,000 twins. In parallel, a subset of 861 individuals participated in eight in-person testing (IPT) sessions, encompassing health, cognition, and memory assessments. Notably, IPT9 and IPT10 introduced daily tracking of memory, emotions, and social interactions, providing valuable insights into short-term fluctuations and early indicators of declining health.

A total of 1,122 blood samples were collected at five time points spanning from 1992 to 2012. Across these five longitudinal waves, participant counts with 1 to 5 measurements were 99, 86, 90, 80, and 30, respectively. Phenotype data, encompassing chronological age, sex, and zygosity, were gathered through comprehensive questionnaires and physical testing at each sampling wave. Following the manufacturer’s protocol optimized for Illumina’s Infinium 450K assay, 200 ng of DNA was prepared for each sample. The bisulfite-converted DNA samples were then hybridized to the Infinium HumanMethylation450 BeadChips using Illumina’s Infinium HD protocol. This process allowed the measurement of DNA methylation levels for 485,512 CpGs in each sample. Following the quality control procedures described by Peters et al., 1,011 samples from 385 twins were retained after processing and quality control on the methylation data (Marjolein Peters et al., 2015). The data is available at ArrayExpress public repository under the accession E-MTAB-7309.

#### Brisbane Systems Genetics Study (BSGS)

Participants BSGS were enrolled in the Brisbane Twin Nevus (BTN) and MAPS (Nevus and cognition studies) projects. The study was approved by the Queensland Institute for Medical Research-Human Research Ethics Committee. Over the span of 16 years, adolescent MZ and DZ twins, alongside their siblings and parents, were recruited for an ongoing investigation into genetic and environmental influences on pigmented nevi and the associated risk of skin cancer and cognition. The study comprises individuals of mainly Anglo-Celtic descent with northern European origins (Joseph Powell et al., 2012).

DNA methylation analysis was conducted on 614 individuals from 117 European descent families recruited through BSGS5. Bisulfite converted DNA samples were hybridized to 12-sample Illumina HumanMethylation450 BeadChips using the Infinium HD Methylation protocol and Tecan robotics, assessing methylation status across 485,577 CpG sites, covering 99% of RefSeq genes. Post-methylation probe filtration by Powell et al. resulted in 417,069 probes. The methylation data are accessible in the GEO public repository under the accessions GSE56105.

#### Genome-wide Methylation Profiles Reveal Quantitative Views of Human Aging Rates (GMPWAR)

This study aims to construct an aging model that gauges one’s methylome age acceleration. Approved by the University of California, San Diego, the University of Southern California, and West China Hospital, the research involved 426 Caucasian and 230 Hispanic individuals, spanning ages 19 to 101, providing extensive methylome-wide profiles. Whole blood samples were collected, and genomic DNA underwent analysis using the Illumina Infinium HumanMethylation450 BeadChip assay, assessing methylation states across 485,577 CpG markers. Stringent quality controls by Hannum et al. ensured the exclusion of unreliable markers and samples (Gregory Hannum et al., 2013). Access to the complete methylation profiles is available at the GEO repository (GSE40279).

#### ScRNA-seq of PMBCs of healthy aging populations

Approval for all human studies was granted by the Washington University in St. Louis School of Medicine Institutional Review Board (IRB-201804084). The study enrolled healthy Caucasian individuals, both males and females, who were non-obese with a BMI below 30, between 2018 and 2019. The research involved specific age and BMI criteria: young males aged 25-29 years (BMI 21.2-28.9 kg/m2, n = 14), and older non-frail males aged 62-70 years (BMI 17.8-29.8 kg/m2, n = 14). Exclusion criteria comprised individuals with a history of cancer, inflammatory conditions, bloodborne diseases, smokers, or recent illness (Denis Mogilenko et al., 2021).

Blood samples (∼100 ml) were obtained via venous puncture in the morning (7–10 a.m.) after an overnight fast. Single-cell suspensions were isolated and underwent droplet-based massively parallel single-cell RNA sequencing using the Chromium Single Cell 5 Reagent Kit according to the manufacturer’s instructions (10x Genomics). Processing involved sample demultiplexing, barcode handling, and cell counting using the Cell Ranger Single-Cell Software Suite (10x Genomics). Subsequent analysis utilized the Seurat R package (Butler et al., 2018), which identified 25 clusters representing primary immune cell populations. Notably, one sample (D18) was excluded from analysis due to its outlier status, identified by distinct clustering in the single-cell RNA sequencing analysis. Raw and processed scATAC-seq data are available at https://www.synapse.org repository (syn22255433)

#### ScRNA-seq of PBMCs of supercentenarians

All experiments using human samples in this study were approved by the Keio University School of Medicine Ethics Committee (approval no. 20021020) and the ethical review committee of RIKEN (approval no. H28-6). PBMCs derived from 7 supercentenarians (SC1–SC7) and 5 controls (CT1–CT5, aged in their 50s to 80s) were profiled using droplet-based single-cell RNA sequencing technology (10× Genomics). Fresh WB from supercentenarians, their offspring residing with them, and unrelated donors was collected, and PBMCs were isolated from WB. Single-cell libraries were prepared from freshly isolated PBMCs by using Chromium Single Cell 3ʹ v2 Reagent Kits. The analysis pipelines in Cell Ranger version 2.1.0 and Cell Ranger R Kit (version 2.0.0) were used for sequencing data processing (Kosuke Hashimoto et al., 2019). Raw UMI counts and normalized expression values for scRNA-Seq are publicly available at http://gerg.gsc.riken.jp/SC2018/.

